# Zika viruses encode multiple upstream open reading frames in the 5′ viral region with a role in neurotropism

**DOI:** 10.1101/112904

**Authors:** Charlotte Lefèvre, Georgia M. Cook, Adam M. Dinan, Shiho Torii, Hazel Stewart, George Gibbons, Alex S. Nicholson, Liliana Echavarría-Consuegra, Luke W. Meredith, Valeria Lulla, Julia C. Kenyon, Ian Goodfellow, Janet E. Deane, Stephen C. Graham, Andras Lakatos, Louis Lambrechts, Ian Brierley, Nerea Irigoyen

## Abstract

Zika virus (ZIKV) is an emerging mosquito-borne flavivirus recently associated with congenital diseases and neurological complications. As for all flaviviruses, the ZIKV RNA genome is expected to encode a single polyprotein with all the enzymatic activities required for viral replication. Here, we report the discovery of multiple non-canonical open reading frames (ORFs) identified by ribosome profiling. In both mammalian and insect cells infected with Asian/American and African ZIKV strains, we observed translation of previously unrecognised upstream ORFs (uORFs) in the 5′ region. In the Asian/American ZIKV lineage, ribosomes translated uORF1 and uORF2 that initiated from non-AUG start codons, whereas in the African ZIKV lineage, these two uORFs were fused into a single uORF (African uORF). Using a reverse genetics system, we examined the impact on ZIKV fitness of the expression of single or dual uORFs by analysing a panel of mutant viruses. We found that expression of the African uORF, and more significantly, the Asian/American uORF1, modulated virus growth and tropism in human cortical neurons and 3D organoid tissue, indicating that these novel uORFs contribute to ZIKV neurotropism. Although ZIKV uORFs are expressed in mosquito cells, they did not have a detectable effect on transmission by the mosquito vector *in vivo*. Our discovery of ZIKV uORFs sheds new light on ZIKV-induced neuropathogenesis and raises the question of their existence in other neurotropic flaviviruses.

## Introduction

Zika virus (ZIKV) is an emerging *Aedes* mosquito-borne *Flavivirus*, a genus which also includes dengue, yellow fever, Japanese encephalitis, tick-borne encephalitis and West Nile viruses. ZIKV was isolated initially from a febrile monkey in Uganda in 1947^1^ but was considered of low importance as most human infections appeared to be asymptomatic. However, it gained prominence from 2007, when the first large epidemic was reported from Yap Island in Micronesia and spread to French Polynesia in 2013^2^. In 2015, the virus reached Brazil and spread across the Americas. Typically, symptoms associated with ZIKV are mild and might include fever, maculopapular rash, conjunctivitis, and myalgia. In adults, the virus can cause severe neurological symptoms, including Guillain-Barré syndrome^2^; and infection of pregnant women can lead to vertical transmission of the virus to cells of the developing brain of the foetus^3^. A key pathological aspect of such infection is impaired development of the unborn brain, leading to gestational abnormalities, including congenital Zika syndrome (CZS). The most well-known feature of CZS is microcephaly^4^, but the syndrome also includes general neurological impairment, neurosensory alterations, delays in motor acquisition and an 11-fold greater risk of death during the first three years of life^5,6^. The devastating impact of ZIKV on newborns has far-reaching social consequences, including stigmatisation of affected mothers and babies plus an increase in illegal abortions across Latin America^7^. Recent epidemiolocal studies also indicate that immunity to ZIKV may not persist for as long as previously thought, with neutralizing antibody levels declining over time in adults^8^.

The relative pathogenesis of African and Asian/American ZIKV isolates has been the subject of debate. The American strain is generally considered more pathogenic, although more serious disease manifestations of the African strain may have been missed in the past because, paradoxically, they were more severe (i.e., early abortion instead of birth defects) and thus, less visible^9,10^. Here, we investigate potential pathological correlates of virus gene expression in representative strains of the Asian/American and African ZIKV lineages in mammalian and mosquito cells through RNA sequencing (RNA-Seq) and ribosome profiling (Ribo-Seq). Like other flaviviruses, ZIKV has a positive-sense, single-stranded RNA genome (gRNA) of ~11 kb, which contains a single long open reading frame (ORF) flanked by 5′ and 3′ untranslated regions (UTRs) of approximately 100 and 400 nucleotides, respectively. The ORF encodes a large polyprotein that is cleaved by host and viral proteases to yield three structural proteins derived from the N-terminal region (capsid - C, precursor/membrane - pr/M, and envelope - E) and seven non-structural proteins (NS1, NS2A, NS2B, NS3, NS4A, NS4B, NS5)^11^. We show here that in addition to the polyprotein, ZIKV encodes previously unrecognised upstream ORFs (uORFs) in the 5′ UTR, initiating from non-AUG start codons. In African ZIKV isolates, a single uORF (African uORF) is present, whereas, in Asian/American strains, this uORF is split into two uORFs (uORF1 and uORF2) by insertion of a single nucleotide.

The impact on ZIKV fitness of the expression of single or dual uORFs is explored by analysing a panel of mutant viruses. We find that expression of the African uORF, and the American uORF1, can modulate virus growth, virulence and tropism in the human brain. We also show that expression of uORF1 is essential for ZIKV neurotropism in the adult and developing brain but has a limited role in the mosquito vector. These experiments reveal novel players in ZIKV pathogenesis and identify new targets for antiviral intervention.

## Results

### Ribosome profiling reveals the presence of novel uORFs in the 5′ UTR of different ZIKV strains

We utilized Ribo-Seq, in combination with whole transcriptome sequencing (RNA-Seq), to investigate translation of the ZIKV genome at sub-codon resolution. Ribo-Seq exploits the capacity of elongating ribosomes to protect ~30 nucleotides of mRNA from digestion during nuclease incubation of cell extracts^12^. Such ribosome-protected fragments (RPFs) are purified and deep-sequenced, revealing the location of translating ribosomes in a cell at the time of harvesting with single-nucleotide precision^13^. African green monkey (Vero) cells and human glioblastoma-astrocytoma (U251) cells were infected at a multiplicity of infection (MOI) of three with the ZIKV American isolate PE243, representative of the Asian/American lineage^14^, and the ZIKV African isolate Dak84 as a model for the original East African strain^15^. To preserve the positions of translating ribosomes upon cell lysis, infected cells at 24 hours post-infection (h p.i.) were flash-frozen or pre-treated for three minutes with the translation inhibitor cycloheximide (CHX) before harvesting. CHX treatment is widely used in Ribo-Seq studies but can lead to the accumulation of 80S ribosomes on start codons and, in stressed cells, can induce the accumulation of RPFs in the 5′ region of coding sequences^16^. For this reason, cells were harvested by flash-freezing to avoid these potential biases (unless stated). Ribo-Seq and RNA-Seq libraries were prepared and deep sequenced as previously described^17^.

At 24 h p.i., the viral envelope (E) protein of each strain is robustly expressed in infected Vero and U251 cells (**Fig 1A**). The Ribo-Seq (red) and RNA-Seq (green) read densities on the virus genome for infected Vero and U251 cells are illustrated in **Fig 1B and Supp Fig 1A-C**. Consistent with the translation of a single polyprotein from the genomic mRNA, read coverage across the main ORF was even, although localized variations in RPF density appear, that may arise from technical biases (ligation, PCR and nuclease biases^18^) or, potentially, from ribosome pausing during translation^19^. Few RPFs were present in the 3′ UTR, consistent with the absence of translation in this region. The 3′ UTR RNA-Seq density was noticeably higher (65-70%) than across the upstream part of the genome in each cell line (**Fig 1B**). Structured flavivirus 3′ UTRs resist degradation by the 5′-3′ Xrn1 host exonuclease, giving rise to noncoding subgenomic flavivirus RNAs (sfRNAs) that accumulate during infection^20^ and are linked to cytopathic and pathologic effects^14^. A sharp spike of RNA-Seq density was observed in the PE243 (nt 10,478) and Dak84 (nt 10,477) libraries (**Supp Fig. 1D and E**), consistent with the presence of a nuclease-resistant RNA structure at this position. Indeed, this location is two nucleotides upstream of the predicted 5′ end of RNA “stem-loop 2” (SL2)^20^.

**Figure 1.**
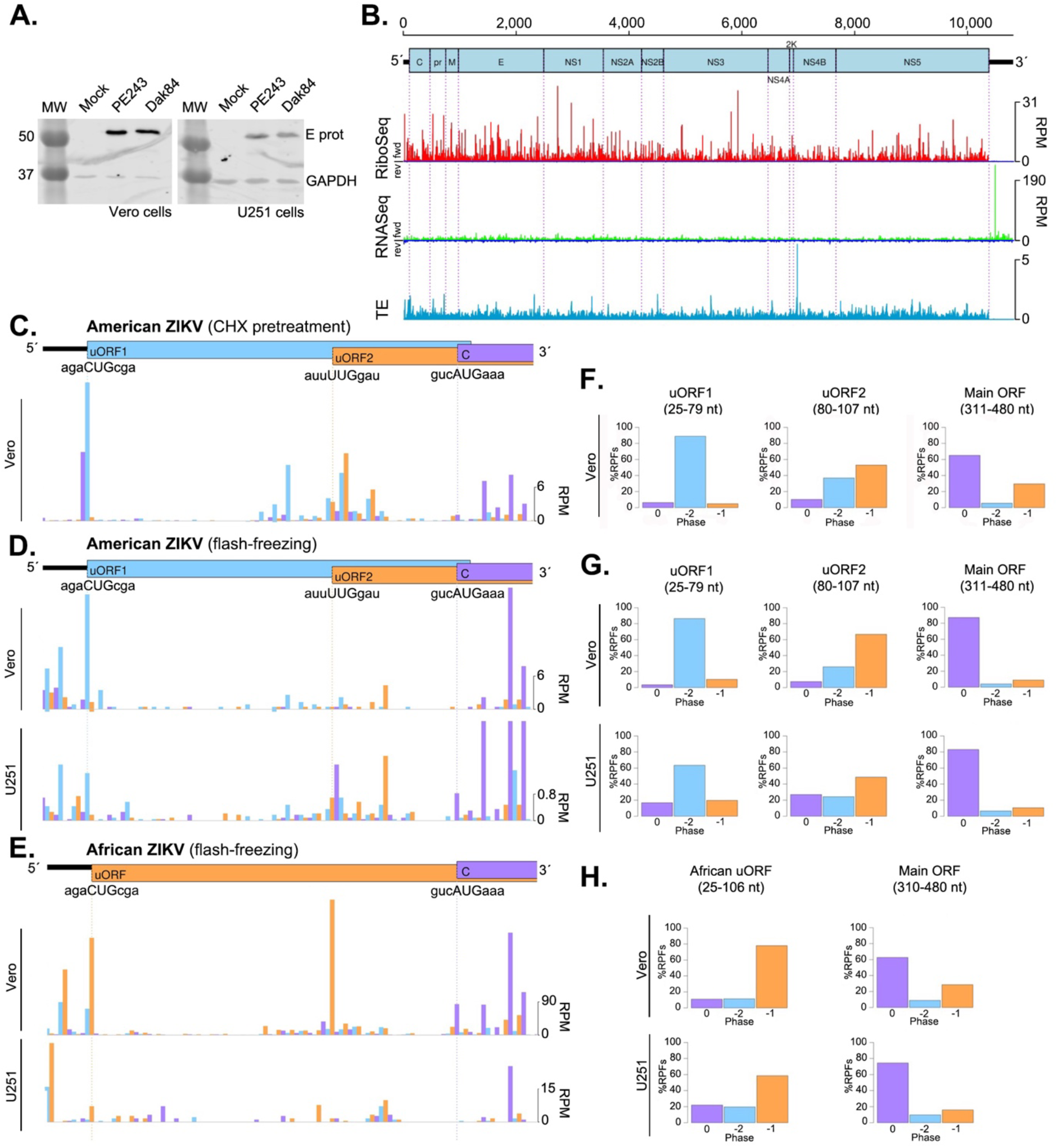
ZIKV RNA synthesis and translation. (**A**) Western blot analysis of ZIKV E protein and GAPDH in Vero and U251 cells infected with PE243 and Dak84 (MOI:3) for 24 h. GAPDH was used as a loading control. Molecular masses (kDa) are indicated on the left. (**B**) Map of the 10807-nt ZIKV/Brazil/PE243/2015 genome. The 5′ and 3′ UTRs are in black, and the polyprotein ORF is in pale blue with subdivisions showing mature cleavage products. Histograms show the read densities, in reads per million mapped reads (RPM), of Ribo-Seq (red) and RNA-Seq (green) reads at 24 h p.i. (repeat 1) in Vero cells pretreated with CHX. The positions of the 5′ ends of reads are plotted with a +12 nt offset to map (for RPFs) approximate P-site positions. Negative-sense reads are shown in dark blue below the horizontal axis. In light blue, the translational efficiency (TE) is calculated as the positive-sense RiboSeq/RNASeq ratio. (**C-D**) The 5′ region of the ZIKV PE243 genome shows two non-AUG uORFs in Vero cells pretreated with CHX (**C**), flash-frozen Vero cells (**D**, upper panel) and flash-frozen U251 cells (**D**, lower panel). (**E**) The 5′ region of the ZIKV Dak84 genome shows the African uORF in flash-frozen Vero (upper panel) or flash-frozen U251 cells (lower panel). Ribo-Seq counts at 24 h p.i. Histograms show the positions of the 5′ ends of reads with a +12 nt offset to map the approximate P-site. Reads whose 5′ ends map to the first, second or third phases relative to codons in the polyprotein reading frame are indicated in blue, orange or purple, respectively. (**F-H**) Bar charts of the percentage of ribosome-protected fragments (RPFs) in each phase relative the polyprotein ORF. Regions with the least amount of overlapping, (coordinates given in plot titles), were selected. Reads whose 5′ ends map to the −2, −1 or 0 phases are indicated in blue, orange or purple, respectively. CHX-treated Vero cells infected with PE243 (**F**); flash-frozen Vero (**G**, upper panel) and U251 (**G**, lower panel) cells infected with PE243; and flash-frozen Vero (**H**, upper panel) and U251 (**H**, lower panel) cells infected with Dak84.

In Ribo-Seq samples, the number of negative-sense reads (dark blue) was negligible (<0.005% of mapped reads), indicating that they are unlikely to be genuine RPFs^13,19^. In comparison, in RNA-Seq samples (**Fig 1B and Supp Fig 1A-C**), a low, uniform coverage of negative-sense reads was observed (at ~1% c.f. positive sense), corresponding to negative-sense intermediates that act as templates for genome replication. The translational efficiency (TE) of each virus genome was calculated by applying a 15-nt running mean filter and dividing the number of Ribo-Seq reads by the number of RNA-Seq reads (light blue), revealing relatively even coverage across the genome. Strikingly, we found substantial TE within the 5′ UTR in both CHX-treated and flash-frozen cells (**Fig 1C and 1D**). In PE243-infected cells, a prominent peak of RPF density was seen at nucleotide 25 of the 5′ UTR in Vero and U251 cells, coinciding with a non-canonical (CUG) initiation codon **(Fig 1C and 1D)**. RPFs mapped in the –2 phase along the length of the associated 29-codon upstream ORF (uORF1, in blue), which ends at nucleotide 111 (the 4^th^ nucleotide of the polyprotein ORF). Additionally, RPFs mapped in the –1 phase to a second uORF (uORF2, in orange), which appears to be translated *via* a non-canonical (UUG) initiation codon at nucleotide 80 (**Fig 1C and 1D**). This uORF2 is 77 codons in length and extends 202 nucleotides into the polyprotein ORF. Analysis of the mapping of Ribo-Seq reads to the ZIKV Dak84 5′ UTR (**Fig 1E**) revealed a single uORF (African uORF), translated in the –1 frame. This African uORF initiates at nucleotide 25 (the same initiation codon used by uORF1) and terminates at nucleotide 309, at the stop codon of uORF2 encoding a putative protein of 94 amino acids in length. In all African ZIKV isolates with available sequence data, uORF1 and uORF2 are present as a single ORF, which appears to have been split in two by the insertion of a uracil residue at position 81 in the 1996 Malaysian lineage that gave rise to the Asian/American strain^21^.

To provide further support for the presence of ZIKV uORFs, we examined the phasing of RPFs in viral 5′ UTRs. For RPFs, mapping of the 5′ end positions to coding sequences (CDSs) characteristically reflects the triplet periodicity (herein referred to as ‘phasing’) of translational decoding^13,17^. We summarized phase relative to the polyprotein (main) ORF by plotting bar charts of the percentage of reads in each phase (**Fig 1F-H**). To avoid the potential confounding effects of overlapping ORFs on these calculations, regions with the least overlap were selected. For uORF1, ribosomal phasing was measured over a 55 nucleotide region (position 25-79), which does not overlap another ORF. Here, a clear dominance of the –2 phase was seen (light blue), consistent with translation in the uORF1 frame (**Fig 1F and 1G**). For uORF2, a short region of 27 nucleotides (position 80-107) was selected, to avoid the high ribosomal occupancy in the 0 phase corresponding to the main ORF that starts at position 108. Here, the majority of reads map to the –1 phase, consistent with uORF2 translation; however, the overlap with uORF1 is still evident from the increased read density in the –2 phase (**Fig 1F and 1G**). For the African uORF (**Fig 1H**), within the chosen region (position 25-106), the majority of reads are attributed to the –1 phase, supporting the expression of this fused uORF. As a positive control, phasing within the polyprotein ORF was assessed, in a region with no known overlapping ORFs (position 310/311-480), revealing a clear dominance of reads attributed to the 0 phase (**Fig 1F-H**). Additionally, as shown in **Supp Fig 2**, the length distribution of Ribo-Seq reads mapping to the ZIKV uORFs mirrored that of polyprotein-mapping RPFs, indicating that they are *bona fide* ribosome footprints.

### The translation of ZIKV uORFs can modulate main ORF expression

The presence of uORFs in mRNAs is often associated with the regulation of downstream gene expression^22,23^. To investigate the potential modulation of ZIKV main ORF expression by the 5′ uORFs, T7 RNA polymerase-capped-derived synthetic reporter mRNAs were prepared in which PE243 uORF1, uORF2 or a 5′ portion of the main ORF was placed upstream of, and in frame with, the firefly luciferase (FF-Luc) reporter gene. The ZIKV and FF-Luc sequences are separated by the short, foot and mouth disease virus 2A autoprotease-encoding sequence that liberates the FF-Luc enzyme following expression in cells (**Fig 2A**). RNAs were reverse transfected into Vero cells alongside a T7-derived RNA expressing Renilla luciferase (Ren-Luc) as transfection control. FF-Luc and Ren-Luc activity was measured at 30 hours post-transfection (h p.t.), and translation efficiencies were determined after normalisation with main ORF (wild-type, WT) translation levels (**Fig 2B**, right panel in red). Under these conditions, uORF1 and uORF2 expression levels were 0.80% and 4.13%, respectively (**Fig 2B**). To assess whether translation of the uORFs could affect main ORF translation, mutations were introduced into uORF1 and uORF2 that were predicted to reduce or increase their expression (**Fig 2B-2D**). As shown in **Fig 2B**, mutation of the uORF1 start codon from CUG to CUA (uORF1-KO, blue) led to a modest reduction in uORF1 expression (left panel), no change in uORF2 expression (middle panel) and a small increase in main ORF translation (right panel). Reducing uORF2 expression by changing the initiation codon from UUG to UUA (uORF2-KO, pink), or introducing a premature stop codon within uORF2 at residue number 6 (uORF2-PTC1, orange), reduced uORF2 translation by 50% (**Fig 2B**, middle panel) and led to a slight decrease in main ORF expression (**Fig 2B**, right panel). Replacing the uORF2 initiation codon with a canonical AUG codon (in purple) led to a substantial increase in uORF2 translation and prevented main ORF expression (**Fig 2B**, middle and right panels). A fusion of uORF1 and uORF2 that recapitulated the African uORF (African-like; in green) did not significantly change uORF expression (**Fig 2B**, middle panel) and led to a modest, albeit significant, increase in main ORF translation (**Fig 2B**, right panel). Similar results were obtained in U251 cells transfected with main ORF-2A-FFLuc mutants (**Supp Fig 3A**).

**Figure 2.**
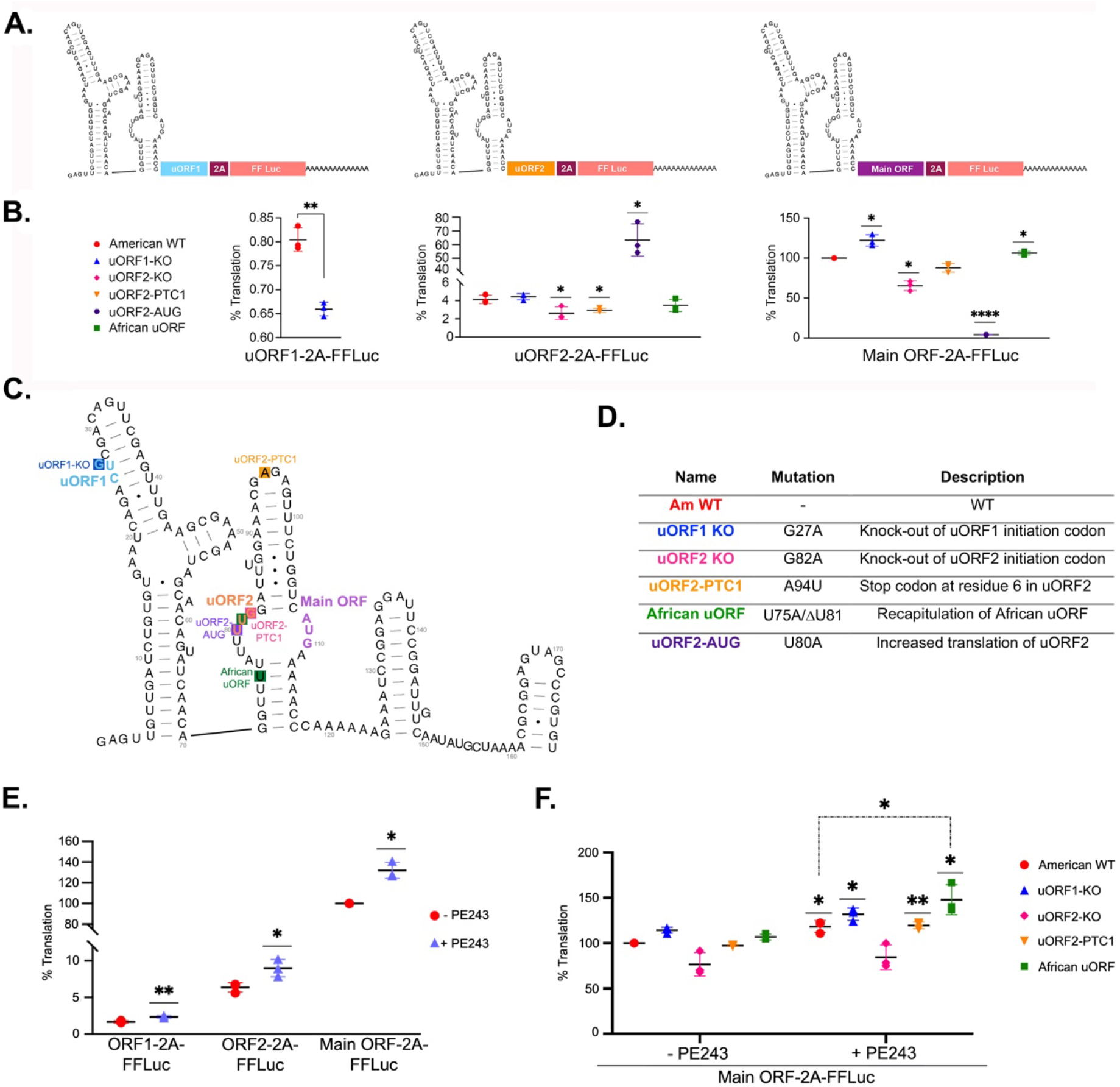
Analysis of ZIKV uORF translation using luciferase reporters. (**A**) Scheme of firefly luciferase (FF-Luc) reporter constructs where the 2A-FFLuc cassette is positioned downstream of and in frame with uORF1 (left), uORF2 (middle) or main ORF (right). (**B**) Relative FF-Luc activity of different mutants for uORF1 (left), uORF2 (middle) and main ORF (right) normalised to Renilla luciferase (Ren-Luc) used as a transfection control. Vero-transfected cells were harvested at 30 hours post-transfection (h p.t.). 100% translation accounted for the main ORF wild-type (WT) translation (right panel in red). (**C**) Scheme of the 5′ region of PE243 with initiation codons for uORF1, uORF2 and main ORF indicated in blue, orange and purple, respectively. Modified nucleotides in the different mutants are indicated by a coloured-filled square associated with the mutant name, briefly described in **D**. (**E**) Relative FF-Luc/Ren-Luc ratio of uORF1-, uORF2-, and main-ORF-2A-FFLuc reporter mRNAs in Vero cells infected with PE243 (MOI:3, purple triangle) or mock-infected (red circle) at 6 h p.t.. Cells were harvested at 24 h p.i.. (**F**) Relative FF-Luc/Ren-Luc ratio of main-ORF-2A-FFLuc mutant reporters in Vero cells infected with PE243 (MOI:3) or mock-infected at 6 h p.t.. Cells were harvested at 24 h p.i.. Experiments were performed in triplicate with three biological replicates. All *t*-tests were two-tailed and did not assume equal variance for the two populations being compared (\**p* < 0.05, ** *p* < 0.01, **** *p* < 0.0001). All *p*-values are from comparisons of the mutant with the respective wild-type in the same ORF.

We went on to ask whether viral infection could influence the relative utilisation of main and uORFs. In these experiments, reporter mRNA-transfected cells were infected at 6 h p.t with PE243 (MOI:3) and harvested 24 h later. In the context of infection, the expression of the uORFs and main ORF relative to each other remained similar, although the total expression of each ORF significantly increased compared to uninfected cells (**Fig 2E**). Notably, the raw luciferase values (**Supp Table 1**) were slightly lower in the presence of the virus, which may reflect some impairment of translation initiation as a result of the phosphorylation of the alpha subunit of the initiation factor 2 (p-eIF2α) during infection (**Supp Fig. 3B**). The effect of uORF mutations on main ORF expression in infected cells was also tested (**Fig 2F**). In all cases, except uORF2-KO, the relative expression of the main ORF was increased modestly. In conclusion, virus infection modestly and uniformly increased expression from upstream and main ORFs.

### uORF translation modulates virus replication

To investigate the potential role of ZIKV uORFs in virus replication, a panel of viruses containing the mutations tested above was generated using reverse genetics of an American ZIKV infectious clone^24^, detailed in **Fig 3A**. Given that structured RNA elements and long-range interactions in the 5′ and 3′ terminal regions of the ZIKV genome are essential for virus translation and replication^25,26^, we began by confirming that the 5′ end structures were retained in full-length RNA transcripts of the mutant viruses. Using selective 2′-hydroxyl acylation analysed by primer extension (SHAPE), we found that the structure of the 5′ UTR and the start of the main ORF of the mutant viruses generally very closely matched that of the American WT infectious clone (**Fig 3B and Supp Fig 4A**). Two differences were observed; modelling of the African-like mutant virus indicated a slightly shorter 3^rd^ stem-loop (cHP), with loss of two base pairs at the bottom of the helix (**Fig 3B**), and in the uORF2-AUG mutant, the internal loop in the centre of the 2^nd^ stem-loop (SLB, **Supp Fig 4A**) is base-paired. Mutant viruses were also analysed for the stability of the introduced mutations. After five passages, RT-PCR analysis of intracellular viral RNA (i.e., initially infected with MOI 0.01 PFU/cell) revealed that all mutations were stable, with two exceptions. ZIKV uORF2-KO reverted to WT after passage 1, and we could not recover any virus following electroporation of the ZIKV uORF2-AUG mutant (**Supp Fig 4B**). These observations suggest that the translation efficiency of uORF2 might be critical to ZIKV replication.

**Figure 3.**
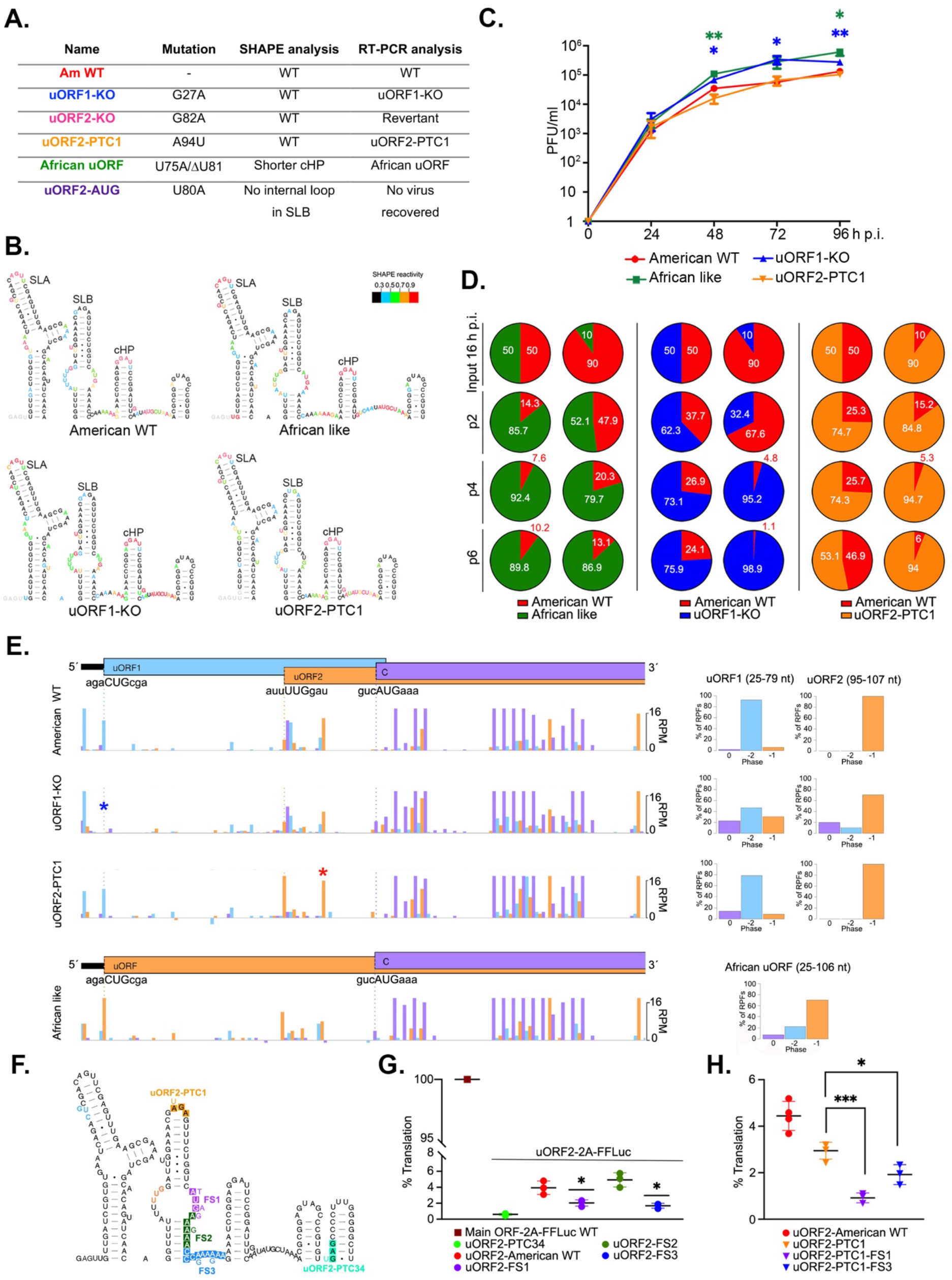
The significance of uORF translation in virus infection. (**A**) Summarising table of SHAPE and RT-PCR data for ZIKV uORFs mutant viruses derived from the pCCI-SP6-American ZIKV infectious clone. (**B**) SHAPE RNA secondary structure of the 5′ region (first 180 nucleotides) of the American WT, African-like, uORF1-KO and uORF2-PTC mutant viruses. Nucleotides are colour-coded based on SHAPE reactivity. SLA (stem-loop A), SLB (stem-loop B) and cHP (capsid hairpin). (**C**) Time-course of infection of U251 cells with ZIKV mutant viruses (MOI 0.01) for 96 h. Plaque assays were performed on serial dilutions of the supernatant containing released virions. Values show the mean averages of the titration of three biological replicates. Error bars represent standard errors. PFU: plaque forming units. All *t*-tests were two-tailed and did not assume equal variance for the two populations being compared (\**p* < 0.05, ** *p* < 0.01). All *p*-values are from comparisons of the mutant virus with the American WT. (**D**) Pie charts of competition assays of American WT with different mutant viruses after 2 (p2), 4 (p4) and 6 (p6) sequential passages in U251 cells. Different proportions of each virus (50:50 and 90:10) were added as indicated in the first row (input 16 h p.i., passage 0). The chart area indicates the RNA proportion for each virus, as measured from sequencing chromatograms of RT-PCR products. Experiments were repeated independently eight times (raw data in **Supp Table 5**). (**E, left panels**) Ribo-Seq read density in the 5′ region of the ZIKV genome at 24 h p.i. of flash-frozen Vero cells infected with American WT, uORF1-KO, uORF2-PTC1 and African-like viruses (MOI:3). Histograms show the positions of the 5′ ends of reads with a +12 nt offset to map the approximate P-site as described in **Fig 1C-E**. Note that in order to properly visualise RPFs across the uORFs, the y-axis has been truncated at 20 reads per million virus mapping reads for the Ribo-Seq samples, leaving some RPF counts for the main ORF off-scale. The blue asterisk indicates the initiation codon of uORF1 and the red asterisk is the premature termination codon in uORF2. (**E, right panels**) Bar charts of the percentage of RPFs in each phase, relative to the main ORF (plot as described in **Fig 1F-H**). (**F**) Scheme of the 5′ region of the American WT. Premature termination codons (PTC) are colour-squared in orange and bright green for uORF2-PTC1 and uORF2-PTC34, respectively. The nucleotides corresponding to the main ORF initiation codon (FS1) and two potential slippery sequences (FS2 and FS3) are coloured in purple, green and blue, respectively. Mutated nucleotides are colour-coded and located adjacent to the original one. Relative FF-Luc activity of different mutants (PTC and FS) of uORF2-2A-FFLuc (**G**) and uORF2-PTC1-2A-FFLuc (**H**) in Vero-transfected cells as in Fig 2B. 100% translation accounted for the main ORF WT (brown square). All *t*-tests were two-tailed and did not assume equal variance for the two populations being compared (\**p* < 0.05, *** *p* < 0.001).

To assess the growth and infectivity of the stable mutants, U251 cells were infected with sequence-verified American WT, African-like, uORF1-KO or uORF2-PTC1 viruses at low multiplicity (0.01 PFU/cell) in a multi-step growth experiment from 0 to 96 h p.i. As shown in **Fig 3C**, from 48 h p.i. onwards, African-like and uORF1-KO mutant viruses reached significantly higher titres (~6-fold) than the American WT, whereas no difference was observed with the uORF2-PTC1 mutant. This phenotype was confirmed in competition assays in which cells were simultaneously infected with a set ratio (50:50 and 90:10) of American WT virus to the corresponding mutant virus to a final MOI of 0.01 PFU/cell (**Fig 3D**). Note that the uORF2-PTC1:American WT comparison was 90:10 instead. Supernatants containing virus particles were harvested 72 h later and used to infect fresh cells at a dilution of 1:200. This regimen was repeated six times^27^, and intracellular RNA was harvested, reverse-transcribed, and Sanger sequenced. Passage 0 corresponds to 16 h post-viral RNA input and represents the starting point. From passage 2, the African-like and uORF1-KO mutant viruses began to dominate the cultures (**Fig 3D**), indicating that both mutant viruses can outcompete American WT. In contrast, the proportion of uORF2-PTC1 viral RNA was relatively unchanged throughout the course of the experiment (**Fig 3D, right panel**), indicating no competition with the American WT virus. The phenotypic differences amongst viruses were also evident, albeit subtler, in Vero-infected cells (**Supp Fig 4C and Supp Fig 4D**).

In an effort to detect expression of uORF products in infected cells, antibodies to expressed proteins and peptides were prepared, but they lacked sensitivity and specificity. Therefore, we returned to ribosome profiling to examine translation of the uORFs in ZIKV mutant viruses (**Supp Fig 4E and Supp Fig 4F**). As seen in **Fig 3E**, reads corresponding to translation of uORF1 (–2 frame) and uORF2 (–1 frame) were visible in the American WT virus, similar to PE243 (**Fig 1C** and **Fig 1D**). The uORF1-KO mutant, as expected, blocked translation of uORF1, with no reads observed at the initiation codon (blue asterisk); and the African-like virus with fused uORFs recapitulated the pattern of uORF translation of the African virus (as in **Fig 1E**). These observations were further confirmed in phasing bar charts, with the highest proportion of reads in the predicted reading frames (**Fig 3E, right panels**). Unexpectedly, reads corresponding to the uORF2 phase were present after the premature termination codon in uORF2-PTC1 (**Fig 3E,** red asterisk), supported by the high proportion of reads corresponding to the –1 phase in phasing plots (**Fig 3E, right panels**). Thus, the translation of uORF2 might also be achieved through an additional mechanism.

### ZIKV uORF2 gets translated by alternative mechanisms

As described earlier (**Fig 2B, middle panel**), knockout of the uORF2 initiation codon or introduction of a premature termination codon (uORF2-PTC1) depressed uORF2 translation only two-fold in comparison to the American WT. In a further attempt to block the synthesis of the uORF2 product, a premature termination codon was placed within the middle of the uORF2 sequence (uORF2-PTC34, **Fig 3F** in bright green) in the context of the uORF2-2A-FFLuc mRNA, and luciferase assays performed in transfected cells. As shown in **Fig 3G**, introduction of PTC34 reduced uORF2 translation ~10-fold, suggesting that a signal for alternative uORF2 expression is present between residue 6 (PTC1) and residue 34 (PTC34). To investigate this, we looked for potential initiation sites between codons 6 and 34. No AUG codons are present, but four potential sites of non-AUG translation initiation could be accessed through leaky scanning of 40S initiation complexes on the mRNA. These amino acids were sequentially mutated (Mut1-AUC, Mut2-AGG, Mut3-AUU and Mut4-ACG; **Supp Fig 4G**) within the uORF2-2A-FFLuc reporter mRNA and tested in transfected cells (**Supp Fig 4H**). In all cases, luciferase levels were unchanged, arguing against a role for leaky scanning in uORF2 expression. We noticed that the sequence immediately downstream of the main ORF AUG has two A-rich stretches upstream of a stem-loop (cHP) that could potentially act as signals for programmed ribosomal frameshifting (PRF^28^), diverting a proportion of ribosomes into the overlapping –1 (uORF2) reading frame (denoted FS2 and FS3 in **Fig 3F**). Reducing the homopolymeric nature of each of these stretches by the introduction of two G residues within the uORF2-2A-FFLuc reporter reduced uORF2 translation at the FS3 site by 50% suggesting that at least half of uORF2 expression is derived from ribosomes that undergo –1 PRF very early during main ORF translation (**Fig 3G**). Disruption of the main ORF initiation codon (FS1) also significantly reduced uORF2 expression, consistent with a requirement for translation of the main ORF in uORF2 expression and frameshifting into the –1 frame at the ‘blue’ slippery sequence (FS3 region). Furthermore, within the context of the uORF2-PTC1-2A-FFLuc reporter mRNA, both FS1 and FS3 mutations reduced uORF2 expression further, presumably as ribosomes can no longer access uORF2 following initiation at the main ORF AUG codon (FS1) and frameshifting into uORF2 at the FS3 signal (**Fig 3H**).

### ZIKV uORF1-encoded protein helps in the formation of the cytoskeletal cage during infection

Bioinformatic analyses suggest no homology of ZIKV uORFs to known proteins. Given their predicted sizes (uORF1, 3.1 kDa; uORF2, 8.4 kDa; and African uORF, 10.3 kDa), however, it is likely that some, or all, will have intrinsic biological activity. To examine their cellular localization, we transiently expressed mCherry-tagged and TAP (Strep-Strep-FLAG)-/FLAG-tagged uORF variants in uninfected and infected mammalian cells, with PE243 and Dak84, and performed subcellular fractionation (**Fig 4A and Supp Fig 5**). As observed in **Supp Fig 5C and Supp Fig 5D**, uORF2 and African uORF were mainly present in the cytoplasm, and there was no relocalization upon infection. However, uORF1, tagged with mCherry or TAP, and in two different cell lines (Vero and U251 cells), appeared not only in the cytoplasm but also in the cytoskeletal fraction (**Fig 4A and Supp Fig 5E-5H**). Note that uORF1-TAP also appeared in the nuclear fraction, but this was probably due to passive diffusion through the nuclear membrane of a small protein. To discern the specific cytoskeletal target of uORF1, we performed confocal analysis with different proteins marking the three types of cytoskeletal filaments: actin for microfilaments, tubulin for microtubules and vimentin for intermediate filaments (**Fig 4B**). It was found that uORF1-TAP could form denser granular structures that co-aligned with an abnormal ‘collapsed’ vimentin (**Fig 4B**, **orange square**), and this was not observed in mock-transfected cells. ZIKV infection reorganises microtubules and intermediate filaments to form a cytoskeletal cage surrounding viral factories^29,30^. These factories are the sites of viral RNA replication and virion assembly, and cytoskeletal remodelling might partly contribute to localizing these processes in a closed environment to prevent access to innate immune defence components. To test specificity, the cytoskeletal phenotype of the uORF1-KO mutant was compared to the American WT. Previously synchronised Vero cells in the G0/G1 phase were infected (at MOI:3) with either virus, the cells fixed at 18 and 24 h p.i. and stained for viral E protein and vimentin (**Fig 4C**). At 18 h p.i., vimentin was perinuclear in all cells, although a denser arrangement was observed in infected cells (**Fig 4C**, upper panel). At 24 h p.i., the cytoskeletal cage, as defined by the collapse of vimentin, was visible in American WT infected cells (**Fig 4C, lower panel orange squares**) but vimentin remained in a perinuclear distribution in uORF1-KO infected cells. We quantified this phenotype by scoring infected cells in four categories; no change in vimentin reorganization (1), increased density of vimentin in the perinuclear region (2), evidence of vimentin collapse (3), and complete collapse of vimentin (4) (see **Supp. Fig 6**). The data derived from scoring between 110-170 cells per condition in two independent experiments are shown in **Fig 4D**. These data indicate that the expression of the uORF1 polypeptide likely helps in the formation of the cytoskeletal cage in infected cells.

**Figure 4.**
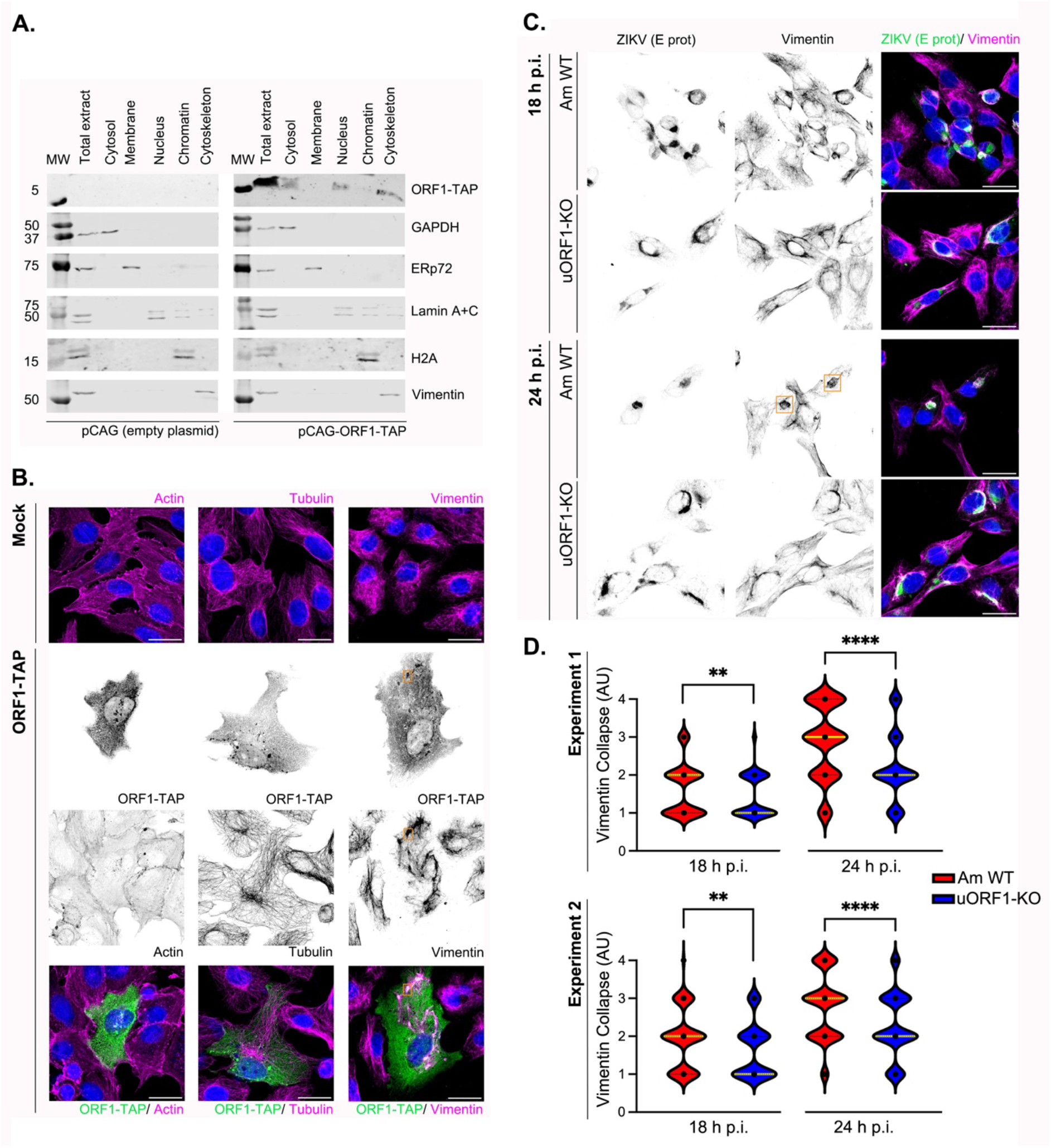
ZIKV uORF1 interacts with the cytoskeleton. (**A**) Subcellular fractionation analysis by Western blot of Vero cells transfected for 48 h with plasmid pCAG (left panel) or pCAG expressing the ZIKV uORF1 protein fused with TAP-tag at the C-terminus (pCAG-ORF1-TAP, right panel). The total extract, cytosolic, membrane, nuclear, chromatin and cytoskeletal fractions were probed with antibodies against FLAG (to detect TAP-tag); GAPDH (cytosolic marker); ERp72 (membrane marker); Lamin A+C (nuclear marker); H2A (chromatin marker); and vimentin (cytoskeletal marker). Molecular masses (kDa) are indicated on the left. (**B**) Representative confocal images of Vero cells transfected for 36 h with pCAG (mock) or the pCAG-ORF1-TAP plasmids. Cells were stained with antibodies against FLAG (green) and different cytoskeletal proteins (i.e. actin, tubulin and vimentin in magenta). Nuclei were counter-stained with DAPI (blue). Images are a maximum projection of a Z-stack. Scale bars, 25 μm. Experiments in **A** and **B** were repeated three times with similar results. (**C**) Representative confocal images of Vero cells infected with the American WT (Am WT) and the uORF1-KO mutant virus (MOI:3) for 18h (upper panel) and 24 h (lower panel). Cells were stained with antibodies against the viral E protein (green) and vimentin (magenta). Nuclei were counter-stained with DAPI (blue). Images are a maximum projection of a Z-stack. Scale bars, 25 μm. (**D**) Violin plots represent vimentin collapse in Vero cells infected with the American WT and the uORF1-KO mutant virus (MOI:3) for 18 and 24 h. Vimentin collapse is quantified in immunofluorescence images as described in **C.** Vimentin collapse units are arbitrary units (AU) which are scored as indicated in the text. 110-170 cells per condition were scored (**Supp Table 6**). Experiments in **C** and **D** were independently repeated two times. All *t*-tests were two-tailed and did not assume equal variance for the two populations being compared (** *p* < 0.01, **** *p* < 0.0001).

### ZIKV uORFs are involved in neurotropism in human cortical neurons and cerebral organoids

To assess whether expression of the different uORFs might influence the neurotropism and neurovirulence of ZIKV, 2D human induced pluripotent stem cell (hiPSC)-derived cortical neuronal mono-cultures and human 3D cortical organoid slice cultures were infected with either American WT, uORF1-KO or the African-like ZIKV.

Differentiated hiPSC-derived glutamatergic cortical neurons (i^3^Neurons^31^) were infected at high MOI (10 PFU/cell based on titre in Vero cells) for 4 days. Released virions in the supernatant were quantified and from 48 h p.i. onwards, mutant virus titres were significantly lower, decreasing by 1.5- to 2-log10 at 96 h p.i. (**Fig 5A**), in sharp contrast to their increased replication in U251 cells (**Fig 3C**). This was further confirmed by immunolabelling for ZIKV E protein and confocal microscopy (**Fig 5B** and **Supp Fig 7**), where spread of infection was less obvious with the African-like virus and infection with the uORF1-KO mutant was limited to single infected cells, suggesting little to no viral spread.

**Figure 5.**
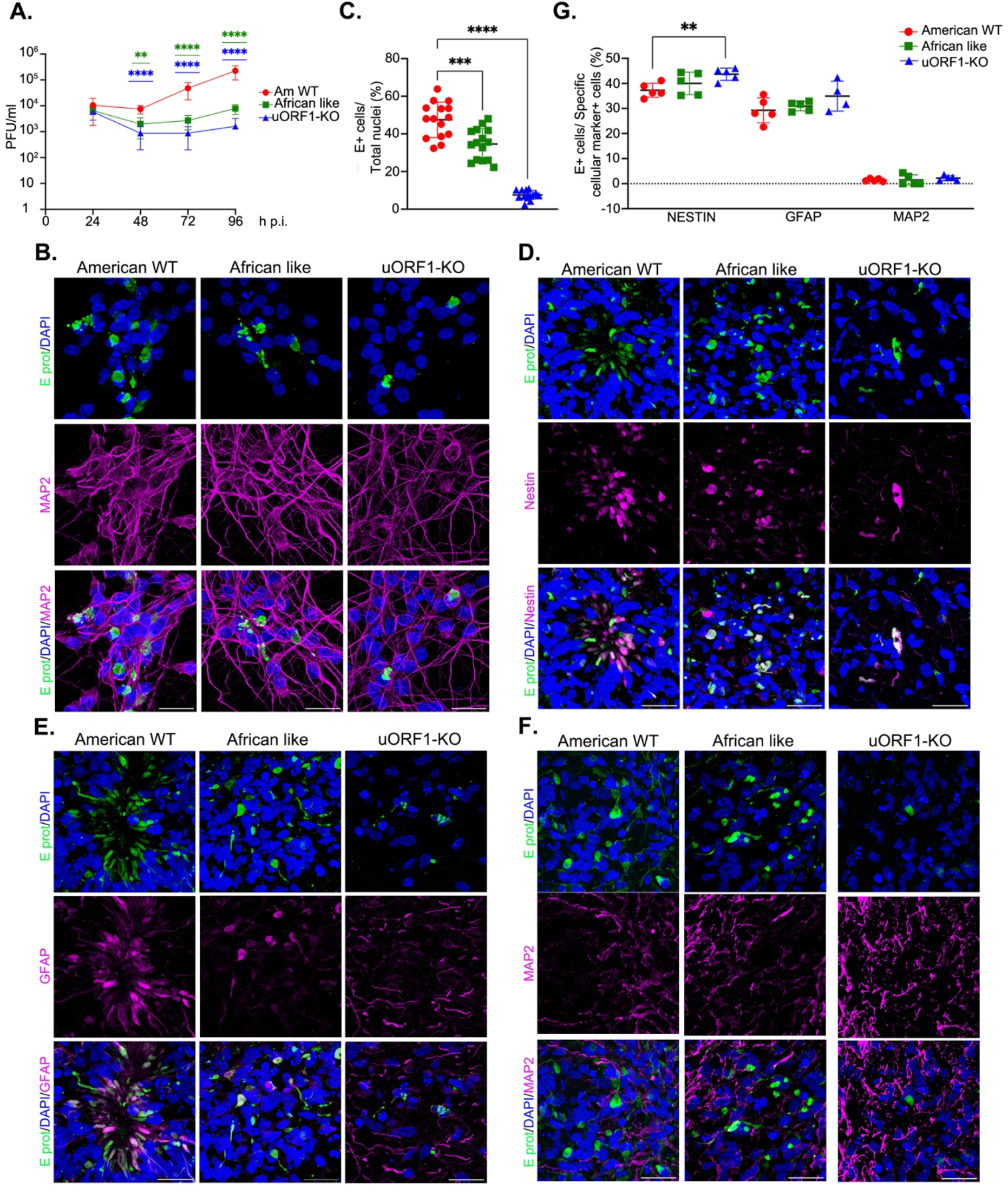
ZIKV uORFs are involved in neurotropism. (**A**) Time-course of i^3^Neurons infected with ZIKV (MOI 10) for 96 h. TCID_50_ were performed with serial dilutions of the supernatant to measure virion release. Values show the mean averages of the titration from five biological replicates. Error bars represent standard errors. PFU: plaque forming units. Statistical analysis was repeated measures two-way ANOVA on the log-transformed data (\*\**p* < 0.01, \*\*\*\**p* < 0.0001). All *p*-values are from comparisons of the mutant virus with the American WT at that specific time-point. (**B**) Representative confocal images of i^3^Neurons infected with the American WT, the African-like and the uORF1-KO viruses (MOI:10) for 96 h. Cells were stained with antibodies against the viral E protein (green) and the mature neuron marker MAP2 (magenta). Nuclei were counter-stained with DAPI (blue). Images represent the maximum projection of a Z-stack. Scale bars, 25 μm. (**C**) Percentage of E^+^ cells in relation to total number of nuclei. 15 images per virus type at 20X resolution (approx. 400-500 nuclei/image) were quantified for E-positive staining. Error bars represent standard errors. Statistical analysis was one-way ANOVA with Gaussian distribution and did not assume equal variance for the two populations being compared (** *p* < 0.01, *** *p* < 0.001, **** *p* < 0.0001). All *p*-values are from comparisons of the mutant virus with the American WT. (**D-F**) Representative confocal images of ALI-COs infected with the American WT, the African-like and the uORF1-KO viruses (MOI:5) for 7 days, showing immunoreactivity for the viral E protein (green) and for different cellular markers (in magenta): nestin (**D**), GFAP (**E**) and MAP2 (**F**). Nuclei were counter-stained with DAPI (blue). Images represent the maximum projection of a Z-stack. Scale bars, 25 μm. (**G**) Percentage of E^+^ cells that are positive for nestin, GFAP and MAP2. 5 images per virus type at 20X resolution (approx. 400-500 nuclei/image) were quantified. Statistical analysis was performed as in **C**.

Next, we used cortical organoids to test the extent of neurotropism in a human brain-like 3D tissue environment and to explore whether uORF1 is also required for the infectivity of other cell-types. To do so, we infected cortical organoid slices grown at the air-liquid interface (ALI-COs), which recapitulate cortical cell type-diversity, layering and neurodevelopmental milestones^32,33^. At 82 days in vitro (DIV), reflecting the first trimester of gestation, ALI-COs were infected with the American WT, the uORF1-KO and the African-like viruses at MOI 5. Virus inoculum was removed after 24 h and the ALI-COs were grown for a further 7 days before being fixed. 6-μm tissue sections were stained for viral E protein and positive cells were quantified in relation to the total number of nuclei. Approximately, 50% of cells were positive for the E protein in the American WT virus infection experiment. This was significantly reduced in the African-like and uORF1-KO virus infection experiments, to ~35% and ~7.5%, respectively (**Fig 5C, Supp Table 2**). This was further confirmed by IF (**Fig 5D-5F, Supp Fig 7**), corroborating our findings on the involvement of the ZIKV 5′ UTR in neurovirulence.

To examine the differential neural cell tropism of these mutant viruses, cryostat sections of ALI-COs were immunostained for a panel of cellular markers, including nestin (as a marker for neural progenitor cells, NPCs), GFAP (a marker for the astroglial lineage such as radial glial cells, glial progenitors and astrocytes), and MAP2 (for mature neurons). As observed in **Fig 5D-G**, ZIKV viruses preferentially infected nestin- or GFAP-positive cells, whereas very few MAP2-positive cells were also positive for E protein. In general, a similar infectivity pattern was seen with all viruses tested (~40% nestin^+^, ~31% GFAP^+^ and ~1.5% MAP2^+^, **Supp Table 3**), suggesting that the modifications to the ZIKV 5′ UTR do not affect neural cell tropism.

Together, these data indicate that the expression of ZIKV uORF1 is an essential neurotrophic factor required for the infection of cortical neurons and NPCs in human tissue. These results also demonstrate that the African 5′ UTR arrangement impairs the infection of NPCs and precursors of the astroglial lineage in the developing brain, although to a lesser extent.

### No detectable effect of ZIKV uORFs in the mosquito vector

ZIKV typically cycles between humans and *Aedes* mosquitoes; hence we also tested whether the uORFs were expressed during the infection of cells derived from the mosquito vector. Ribo-Seq analysis of *Aedes albopictus* (C6/36) cells infected with PE243 for 24 h suggested that both uORFs were occupied by ribosomes during infection (**Fig 6A**). We compared the transmission dynamics of American WT, African-like and uORF1-KO viruses in female *Aedes aegypti*. Mosquitoes were exposed to an artificial, infectious blood meal containing an expected virus titre of 2 x 10^6^ PFU/mL of blood. Actual titres ranged from 0.8 x 10^6^ to 3.6 x 10^6^ PFU/mL across the different experiments, and this uncontrolled variation was accounted for in the statistical analysis. At several time points after the infectious blood meal, we determined the rates of mosquito infection and systemic viral dissemination by RT-PCR and detected the presence of ZIKV in saliva by infectious assay. We calculated infection prevalence as the proportion of blood-fed mosquitoes with a body infection (determined by RT-PCR, **Fig 6B left panel**), dissemination prevalence as the proportion of infected mosquitoes with a virus-positive head (determined by RT-PCR, **Fig 6B middle panel**) and transmission prevalence as the proportion of mosquitoes with a disseminated infection that had infectious saliva (determined by focus-forming assay, **Fig 6B right panel**), as previously described^34,35^. A total of 99, 128 and 234 individual mosquitoes were analysed in the first, second and third experiments, respectively.

**Figure 6.**
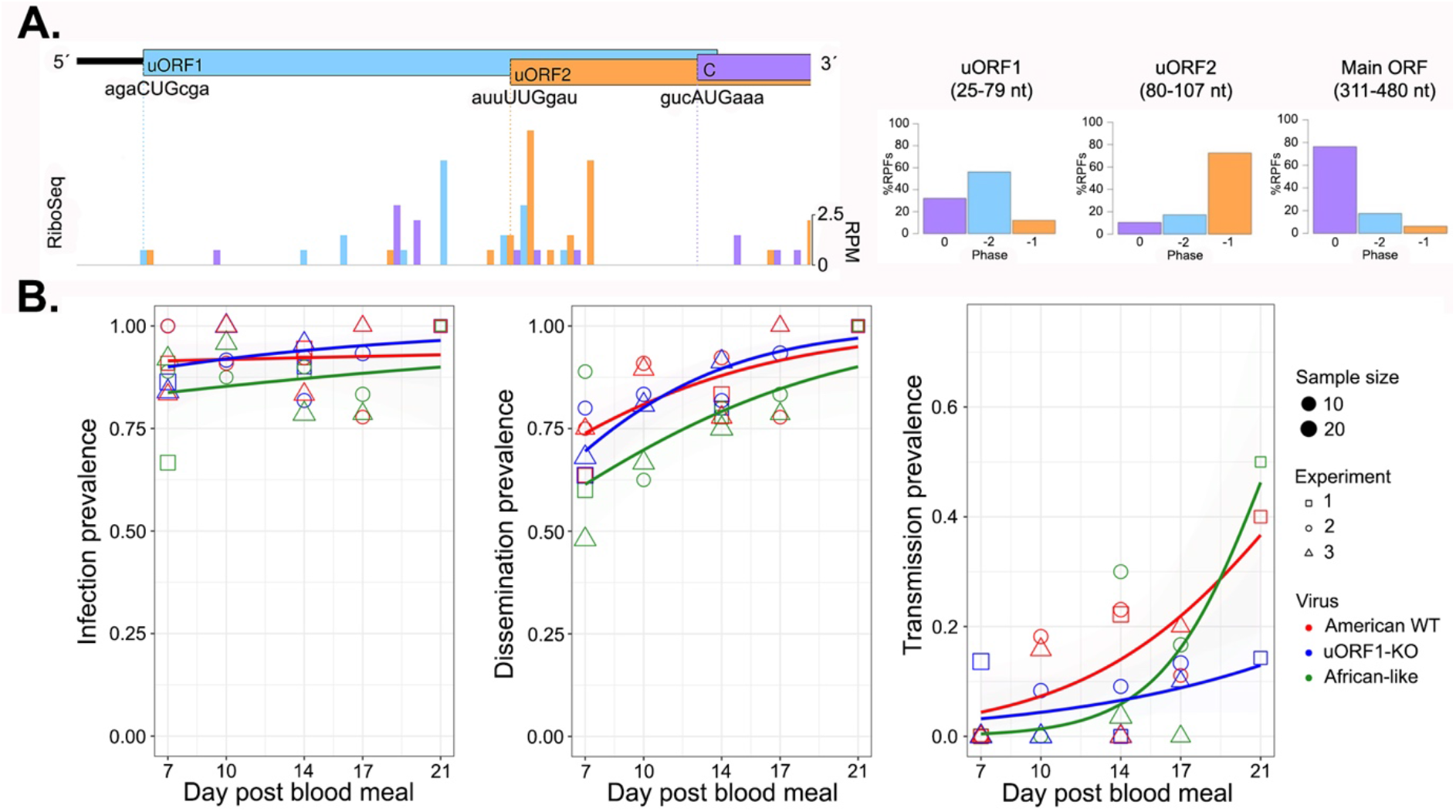
ZIKV mutant viruses display similar transmission dynamics in mosquitoes *in vivo*. (**A, left panel**) Ribo-Seq reads in the 5′ region of the ZIKV genome at 24 h p.i. of CHX-treated C6/36 cells infected with PE243 (MOI:3). Histograms show the positions of the 5′ ends of reads with a +12 nt offset to map the approximate P-site as described in **Fig 1C-E**. (**A, right panel**) Bar charts of the percentage of RPFs translated in each frame in relation to the main ORF as described in **Fig 1F-H**. (**B**) Prevalence of ZIKV infection (**left**), dissemination (**middle**) and transmission (**right**) over time in mosquitoes exposed to an infectious blood meal containing 2 x 10^6^ PFU/mL of virus. Infection prevalence is the proportion of blood-fed mosquitoes with a body infection (determined by RT-PCR), dissemination prevalence is the proportion of infected mosquitoes with a virus-positive head (determined by RT-PCR), and transmission prevalence is the proportion of mosquitoes with a disseminated infection and infectious saliva (determined by focus-forming assay). The data represent three separate experiments depicted by different symbols, colour coded by different ZIKV mutants. The size of the symbols is proportionate to the number of mosquitoes tested, and the lines are the logistic fits of the time effect for each type of virus (ignoring the experiment effect on transmission prevalence in the visual representation).

Most mosquitoes became infected across experiments, different viruses and time points (mean: 90.5%; median: 90.9%; range: 66.7%–100%), and none of these variables had a statistically significant effect on infection prevalence (**Supp Table 4**). Both dissemination and transmission prevalence significantly increased over time (**Fig. 6B-C**), but there was no detectable difference between the different ZIKV mutant viruses (**Supp Table 4**). We conclude that the different ZIKV mutant viruses have similar transmission dynamics in mosquitoes regardless of their 5′-UTR arrangement.

## Discussion

In this study, we provide the first high-resolution maps of ZIKV translation in both mammalian and mosquito cells and describe two previously overlooked uORFs (uORF1 and uORF2) in contemporary ZIKV strains, which exist as a single uORF in African isolates. By generating mutant viruses, we show that the uORFs can modulate virus growth, virulence and tropism in human brain cells. How the uORFs exert their effects is uncertain, but is likely through modulation of expression of the main polyprotein ORF and *via* properties of their encoded proteins.

The translational activity of the ZIKV uORFs is noticeably lower than that of the main polyprotein ORF, consistent with their initiation at non-AUG codons. Whilst the effect of flanking bases on initiator AUG codons is well established (Kozak context), it is less well understood for non-AUG codons, making the strength of non-AUG initiation codons harder to predict^36^. Based on our present understanding, the ZIKV uORFs rank as modest start codons^36,37^. However, their utilisation may also be modulated by RNA structure and, potentially, by virus and host proteins. Recent toeprinting analysis of ZIKV initiation *in vitro* has also detected the utilisation of non-AUG initiation sites at nucleotide positions UUG_41_, UUG_80_ and UUG_86_ (c.f. CUG_25_ and UUG_80_ here). The magnitude of the toeprints observed was influenced by the availability of particular translation factors in the reconstituted assay system and the RNA circularisation status of the mRNAs used^26^. The absence of a toeprint at CUG_25_ in the latter study might reflect the different experimental milieu (*in vitro* reconstitution versus infected cells).

In luciferase reporter gene assays, expression of uORF1 had an inhibitory effect on main ORF translation, whereas expression of uORF2 or the African uORF enhanced translation of the reporter gene. This might explain why mutant viruses bearing these changes (uORF1-KO and African-like) in the 5′ UTR grew to higher titres and were fitter, in comparison to American WT, in tissue culture cells. The inhibitory effect of the uORF1 start codon mutation might be related to the fact that the uORF1 stop codon (UGA) overlaps with the initiation codon of the main ORF (AUGA). We speculate that ribosomes terminating at the stop codon may decrease access of ribosomes to the main ORF AUG, which would be obviated in uORF1-KO, as fewer ribosomes are delivered to the stop codon. In a similar vein, ribosomes translating uORF2 or the African uORF would unwind SLB without terminating within it, increasing access of ribosomes to the main ORF AUG. Viruses with mutations intended to increase (uORF2-AUG) or decrease (uORF2-KO) translation of uORF2 were non-viable or reverted to the wild-type sequence. The dramatic effect of the uORF2-AUG mutation in the virus context could be mediated in several ways. Upregulation of uORF2 expression may drastically reduce the number of scanning ribosomes reaching the main ORF AUG and diminish polyprotein expression, an idea consistent with the significant reduction of main ORF expression in reporter gene assays (**Fig. 2B**). Alternatively, overexpression of the uORF2 protein product in infected cells may be detrimental to virus growth. We also note from SHAPE analysis that this RNA adopts a slightly different fold in the SLB region, which could also contribute to improper replication and translation. The uORF2-KO virus revertant phenotype might be attributable to a reduced amount of uORF2 protein product in infected cells, or altered translational dynamics in the complex 5′ UTR of ZIKV.

Our results, and those of others^38,39^, reveal that eIF2 becomes phosphorylated on its α-subunit upon ZIKV infection (**Supp Fig. 3B**). eIF2α phosphorylation leads to a general shutdown of protein synthesis by inhibiting translation initiation, but in some cases, such as the translation of the activation transcription factor 4 (ATF4^40^), uORFs act as enhancers of main ORF translation as a response to cellular stress. The finding that the ZIKV ORFs are translationally enhanced during infection suggests that the ZIKV 5′ arrangement, and the likely interplay of the two uORFs and main ORF, may offer some resistance to the translation inhibition engendered by phosphorylated eIF2α. This is most evident with the African-like arrangement (**Fig. 2F**) and might explain why the African-like virus outgrows American WT in tissue culture (**Fig. 3D**).

An unexpected finding is the dual mode of uORF2 expression, via initiation from the UUG initiation codon, and also expression of an N-terminally truncated version by –1 ribosomal frameshifting of ribosomes initiating at the main polyprotein (0) frame. Based on previous work^28^, a likely site of frameshifting in this region is the C_CCA_AAA putative “slippery” sequence located four codons into the polyprotein and three nucleotides upstream of the cHP stem-loop. Consistent with this hypothesis, mutation of this sequence to non-slippery C_CCG_AAG reduced uORF2 expression in reporter gene assays (**Fig. 3H**). Frameshifting to CCC_AAA (uORF2 frame) generates an N-terminal variant of uORF2, in which five amino acids derived from the start of the polyprotein replace the 14 N-terminal residues from initiation at the UUG codon. Conceivably, ‘non-canonical’ expression of uORF2 could also arise as a result of transcriptional slippage of T7 RNA-polymerase on the A-rich tract in this region, and indeed, transcriptional slippage has been reported to account for the proposed frameshifting that takes place at a similar position in the hepatitis C virus (HCV) genome^41^. However, analysis of the ZIKV RNA-Seq data does not reveal any heterogeneity in this A-rich region, in contrast to what was observed with HCV^41^.

Bioinformatics analyses do not reveal any apparent homology of ZIKV uORFs to known proteins, although AlphaFold 2 analysis^42^ suggests that uORF1 adopts an alpha-helical conformation, whereas uORF2 is likely to have no well-defined secondary structure (**Supp Fig. 8A**). Related uORFs can be identified in other neurotropic flaviviruses, such as Japanese encephalitis virus (JEV)^43^ and West Nile virus (WNV) (**Supp Fig. 8B**), and they are also predicted to have adopted alpha-helical conformations. The expression in *trans* of uORF2 and the African uORF resulted in a cytoplasmic localization that was not changed upon infection. However, transfected uORF1 was predominantly cytoskeletal and led to the collapse of the vimentin network, with no change observed in microfilaments or microtubules. Previous electron tomography experiments have revealed that ZIKV infection leads to a reorganization of microtubules and intermediate filaments, forming a cytoskeletal cage that surrounds viral factories^29,30^. The remodelling of the cytoskeleton around these sites of viral RNA replication and virion assembly confines these processes to a closed environment where components of the innate immune response cannot sense them. Our results reveal that uORF1-KO viruses are delayed in the formation of the cytoskeletal cage in infected cells and suggest a potential role of uORF1 in avoiding innate immune recognition.

Vimentin is a type III intermediate filament and displays typical features of such proteins, with an N-terminal domain (~100 aa), a central α-helical rod (310 aa) and a C-terminus of variable length (15-1400 aa). The central rod domain allows the formation of coiled-coil dimers that can assemble into higher-order filaments. The bundling of peripheral vimentin into thicker assemblies in the perinuclear area of the cells, as observed upon uORF1 transfection, has been previously observed in normal cells^44^. It has been associated with post-translational modifications of vimentin, such as phosphorylation of the head domain^44,45^ or deacetylation (by histone deacetylase 6) upon expression of certain oncogenes^46^. It is unknown whether vimentin is subjected to any post-translational modifications upon ZIKV infection but, based on the α-helical prediction of AlphaFold2, the uORF1 protein could potentially interact with the central rod domain of vimentin, intercalate during dimer formation, and antagonise correct polymerization.

In addition, vimentin, in conjunction with nestin, are the first intermediate filaments expressed during early brain development^47^. During neural development, the cytoskeleton is involved in different processes (e.g., orientation of the mitotic spindle, interkinetic nuclear migration, microtubule organising centre centrosome activity and assembly of primary cilia) that are fundamental for the correct generation and maturation of neurons^48^. However, the disruption of any of these processes impairs neurogenesis (differentiation of NPCs into mature neurons) in the developing brain, provoking neurological defects such as microcephaly, the key pathological aspect of ZIKV infection. It has been previously described that ZIKV infection can trigger premature differentiation by increasing vertically oriented NPCs^49^ and shortened cilia^50^. Our results suggest that the interaction of uORF1 with the cytoskeleton might also play a role in premature neurogenesis, although this needs further characterisation in a relevant human model that can mimic the 3D organisational structure of the developing brain, such as cerebral organoids.

In this work, we have used ALI-COs, a highly accessible slice culture version of whole-cortical organoids for viral infections, with improved oxygenation, nutrition and longevity^32,33^. This allowed optimal characterisation of the neurotropism exerted by the different ZIKV 5′-UTR mutant viruses. Our finding that the uORF1 knock-out virus has a defect in infecting ALI-COs suggests that the protein encoded by uORF1 - only present in the Asian/American strains - is an essential neurovirulence factor required for ZIKV neuropathogenesis in the developing brain. Interestingly, a virus with the African uORF in the American virus background was also impaired, although to a lesser extent, in brain organoids and iPSC-derived cortical neurons. This may explain, at least in part, how a mutation in the 5′ region in a rarely detected pathogen such as ZIKV drove a major epidemic in the Americas in 2015/2016. Until now, the only mechanism associated with an increase in neuropathogenesis in epidemic Asian/American strains compared to pre-epidemic Asian strains (e.g., Cambodia 2010) was the S139N mutation in the membrane precursor (prM) protein^51^. However, the different mutant viruses had similar infection, dissemination, and transmission dynamics in *Aedes* mosquitoes, regardless of their 5′-UTR arrangement, although these uORFs are also translated in the mosquito vector, suggesting that they may uniquely function in mammalian hosts.

It is, therefore, necessary to further investigate and characterize the role of these 5′ uORFs in the neurovirulence and neuropathogenesis of flaviviruses. To date, they have only been described or predicted in neurotropic flaviviruses (ZIKV, JEV^43^ and WNV, **Supp. Fig. 8B**) and they could drive phenotypic divergence with visceral flaviviruses such as dengue or yellow fever viruses.

## Materials and Methods

### Cells and viruses

Vero (ATCC, CCL81) and U-251 (ATCC, CRL-1620) cells were maintained in Dulbecco’s modification of Eagle’s medium (DMEM) supplemented with 5% and 10% (vol/vol) foetal calf serum (FCS), respectively. Culture medium also contained 100 U/ml penicillin, 100 μg/ml streptomycin and 2 mM L-glutamine. Cells were incubated at 37 °C in the presence of 5% CO_2_. *Aedes albopictus* C6/36 cells (ATCC, CLR 1660) were maintained in L-15 medium supplemented with 10% FCS at 28 °C without CO_2_. The American strain ZIKV/Brazil/PE243/2015 (Genbank accession number KX197192.1) was kindly provided by Prof. Alain Kohl (Centre for Virus Research, University of Glasgow, United Kingdom), and the African strain ZIKV/*A.taylori*-tc/SEN/1984/41662-DAK (Genbank accession number KU955592.1) was obtained from the European Virus Archive. ZIKV mutant viruses were rescued by electroporation of Vero cells with SP6-transcribed RNAs of pCC1-SP6-ZIKV derivatives as previously described^24^. Virus stocks were amplified in Vero cells and concentrated by ultracentrifugation (110,500 × g for 3h at 4°C). All mutant viruses were passaged five times at a low MOI (0.01), and viral RNA was subjected to RT-PCR analysis to confirm the presence or reversion of the introduced mutations.

Upon reaching 70-80% confluence, Vero and U251 cells were infected with parental or mutant viruses in serum-free DMEM supplemented with 100 U/ml penicillin, 100 μg/ml streptomycin and 2 mM L-glutamine at the indicated MOI. After adsorption for 1h at 37 °C, the inoculum was removed and replaced with DMEM containing 2% FCS and 20 mM HEPES pH 7.4. In cases where the infection was preceded by transfection, the transfection medium was washed three times with PBS prior to infection. C6/36 cells were infected as above but using L-15 medium. Mosquito cells were incubated at 28 °C.

### Plasmids

A pCC1BAC vector containing the open reading frame of the ZIKV BeH819015 isolate flanked by the 5′ and 3′ UTRs for ZIKV PE243 (pCC1-SP6-Am ZIKV)^24^ was kindly provided by Prof. Andres Merits (University of Tartu, Tartu, Estonia). Mutations in the 5′ UTR were generated by site-directed mutagenesis with the indicated oligonucleotides **(Supp Table 7)** in a subcloned gene fragment of the first 2,000 nucleotides of ZIKV PE243 maintained in pcDNA.3. Following sequencing to confirm no inadvertent changes had arisen, mutated fragments were cloned back into pCC1-SP6-Am ZIKV at the *EcoR1* and *AvrII* sites.

The previously described pSGDLuc vector^52^ was used as a template to design the Renilla and the Firefly luciferase reporters (pREN-Luc and pFF-Luc). The pREN-Luc cassette was amplified using the following oligonucleotides 5′-TCCGCCCAGTTCCGC CCATTCTCCGC and 5′-GCGCTCTAGATTATCTCGAGGTGTAGAAATAC and cloned back into pSGDLuc previously digested with *AvrII* and *XbaI*. This created a plasmid expressing only the Renilla luciferase gene under the T7 promoter. For the FF-Luc reporters, the first 194 nucleotides of the ZIKV genome were cloned in the pSGD plasmid previously digested with *XhoI* and *BglII* using a different set of oligonucleotides **(Supp Table 7)** to introduce the uORF1, uORF2 and Main ORF in frame with the 2A-Firefly luciferase gene. In addition, a T7 promoter was introduced upstream of the ZIKV coding region. Mutations within the uORFs were introduced by site-directed mutagenesis.

The uORF1, uORF2 and African uORF coding sequences were amplified using specific oligonucleotides **(Supp Table 7)** and cloned in frame at the C-terminus with a FLAG-, TAP- or mCherry tag. *PacI-* and *AflII*-digested PCR fragments were ligated into pCAG previously digested with these restriction enzymes. All sequences were confirmed by Sanger sequencing.

### RNA transcript preparation and luciferase reporter assays

Transcription reactions were performed using the mMESSAGE mMACHINE™ T7 transcription kit (Thermo Fisher Scientific) for 2 h at 37 °C and terminated by treatment with TURBO™ DNase (Thermo Fisher Scientific) for 30 min at 37 °C. Prior to transfection, RNA transcripts were polyadenylated for 30 min at 37 °C. Vero and U251 cells were transfected in triplicate with Lipofectamine 2000 reagent (Invitrogen) by RNA reverse-transfection, where cells in suspension were added directly to the RNA complexes in 96-well plates. For each transfection, 100 ng of purified T7 ZIKV-uORFs FF-Luc RNA and 5 ng of purified T7 Ren-Luc RNA plus 0.3 μl Lipofectamine 2000 in 20 μl Opti-MEM (Gibco-BRL) supplemented with RNaseOUT (Invitrogen; diluted 1:1,000 in Opti-MEM) was added to each well containing 10^5^ cells. RNA to be transfected was purified using an RNA Clean and Concentrator kit (Zymo Research). Transfected cells in DMEM supplemented with 2% FCS and 20 mM HEPES pH 7.4 were incubated at 37 °C for the indicated times. At 6 hours post-transfection, cells were infected with PE243 and Dak84 at MOI:3 by adding 50 μl of DMEM-2% FCS medium-containing virions. Firefly and Renilla luciferase activities were determined using the Dual-Luciferase Stop & Glo Reporter Assay System (Promega). Translation efficiency was calculated as the ratio of FF-Luc units (uORFs-/Main ORF-translation) to Ren-Luc units (transfection control), normalized by the ratio of Main ORF WT-translation.

### DNA transfection

Vero and U251 cells were transiently transfected with pCAG-uORF plasmids using a commercial liposome method (TransIT-LT1, Mirus). Transfection mixtures containing plasmid DNA, serum-free medium (Opti-MEM; Gibco-BRL) and liposomes were set up as recommended by the manufacturer and added dropwise to the tissue culture growth medium.

### Plaque assays

Viral titres were determined by plaque assay. Viral supernatant was 10-fold serially diluted in serum-free DMEM. Vero cells in 6-well plates were infected with 400 µL of diluted supernatant. After 1h at 37°C with regular rocking, the inoculum was removed and replaced with a 1:1 mixture of 3% low-melting agarose and MEM 2X (20% 10X MEM, 8 mM L-glutamine, 0.45% Na_2_CO_3_, 0.4% non-essential amino acids, 4% HEPES pH 7.4 and 4% FCS). Plates were incubated at 37°C for 5 days prior to fixing with formal saline (4% formaldehyde and 0.9% NaCl) at room temperature. Cell monolayers were stained with 0.1% toluidine blue to visualise plaques. The number of plaque-forming units per mL (PFU/mL) was calculated as n° plaques * 2.5. The final number of PFU/mL per time point was calculated by averaging the results of three biological repeats.

### TCID_50_ assays

ZIKV replication was assessed using a 50% tissue culture infective dose (TCID_50_) assay in Vero cells. Supernatant derived from infected i^3^Neurons cells was subjected to 10-fold serial dilutions. Cells were fixed and stained at 96 h p.i. as indicated above. Wells showing any cytopathic effect (CPE) were scored positive.

### Virus competition assays

Dual infection/competition assays were performed in duplicate on Vero and U251 cells using the American WT and different mutant viruses at an equal or 9:1 ratio at a total MOI of 0.1. Media collected from infected plates was used for eight blind passages using 1:10,000 volume of obtained virus stock (corresponding to MOI 0.05–0.2). Total RNA from passages 0 (16 h post-infection to determine viral input), 2, 4 and 6 was isolated as previously described^17^. cDNA was synthesised from 1,000 ng total RNA using M-MLV Reverse Transcriptase (Promega) and 1 µL random primers (1:25 dilution of Random Primers, Invitrogen™). The ZIKV 5′ UTR region was amplified using specific primers **(Supp Table 7)**. Following PCR reactions, the resulting amplicons were subjected to electrophoresis in 3% agarose gels, and DNA bands were purified using the Zymoclean Gel DNA recovery kit (Zymo Research) and subjected to Sanger sequencing. Quantification was performed by integrating area under the curve for each nucleotide and expressing it as percentage of the total signal at that base.

### Subcellular fractionation

Vero or U251 cells seeded in 4-cm dishes were transfected with plasmids of interest. At 40h post-transfection, samples were collected and immediately subjected to subcellular fractionation using the ‘Subcellular Protein Fractionation Kit for Cultured Cells’ (ThermoFisher Scientific). Purified fractions were stored in Laemmli’s buffer at −20°C for downstream biochemical characterisation.

### Immunoblotting

Cells were lysed in 1X Laemmli’s sample buffer. After denaturation at 98 °C for 5 minutes, proteins were separated by 12 % SDS-PAGE and transferred to a 0.45 μm nitrocellulose membrane. For proteins below 16 kDa, 20% SDS-PAGE and 0.22 μm nitrocellulose membrane was used instead. Membranes were blocked (5% non-fat milk powder or bovine serum albumin in PBST [137 mM NaCl, 2.7 mM KCl, 10 mM Na_2_HPO_4_, 1.5 mM KH_2_PO_4_, pH 6.7, and 0.1% Tween 20]) for 30 min at room temperature and probed with specific primary antibodies **(Supp Table 8)** at 4°C overnight. Membranes were incubated in the dark with IRDye-conjugated secondary antibodies diluted to the recommended concentrations in PBST for 1 h at room temperature. Blots were scanned using an Odyssey Infrared Imaging System (Licor).

### Selective 2′ hydroxyl acylation analysis by primer extension (SHAPE)

SHAPE reactions were conducted as previously described^53^. 2 pmol of purified *in vitro* transcribed RNA template (full-length WT ZIKV RNA and mutant derivatives) was denatured at 95 **°**C for 3 minutes, then incubated on ice for 3 minutes before the addition of folding buffer (final concentration 100 mM HEPES pH 8, 6 mM MgCl_2_, 100 mM NaCl). Refolding was conducted at 37 **°**C for 30 minutes. Samples were divided into two equal volumes and treated with either N-methylisatoic anhydride (NMIA) dissolved in DMSO (10 mM final concentration) or DMSO only, for 45 min at 37 °C. Samples were ethanol precipitated and resuspended in 10 μL ddH_2_0 before being reverse transcribed. NMIA or DMSO treated RNA samples were mixed with 5′ NED labelled primer (final concentration 1 μM) (5′ CTTCCTAGCATTGATTATTCTCAGCA TG 3′, Applied Biosystems). Annealing was conducted at 85 **°**C for 3 minutes, 60 **°**C for 10 minutes, and 35 **°**C for 10 minutes. Reverse transcription with Superscript III (Invitrogen) was conducted according to the manufacturer’s instructions at 53 **°**C for 40 minutes. Sequencing ladders were prepared by reverse transcription of 1 pmol of non-treated RNA in the presence of ddCTP, using 5′ VIC labelled primer. All primer extensions were terminated by alkaline hydrolysis, neutralised with HCl and ethanol precipitated before being resuspended in ddH_2_0. The sequencing ladders were divided equally between cDNAs from NMIA- and DMSO-treated RNA. The final samples were dried onto sequencing plates before being resuspended in formamide and fractionated by capillary electrophoresis (Applied Biosystems 3130xl analyser, University of Cambridge Department of Biochemistry DNA Sequencing Facility). SHAPE was performed in triplicate for each genome from separate RNA preparations. Each SHAPE dataset was analysed with the QuSHAPE software^54^. The average normalised NMIA reactivity was calculated for each individual nucleotide and used to model RNA secondary structure using the RNA structure software^55^ using the SHAPE reactivity profile as soft pseudo-free energy constraints. RNA secondary structures were drawn using XRNA (http://rna.ucsc.edu/rnacenter/xrna/xrna.html).

### Immunofluorescence microscopy

Vero or U251 cells were seeded in a 24-well plate with coverslips to be transfected and/or infected. At the indicated times, cells were fixed with 50:50 methanol: acetone for 10 minutes. Coverslips were blocked for 30 minutes in 5% bovine serum albumin (BSA) in 1X PBS at room temperature. Coverslips were incubated in 1% BSA with specific primary antibodies **(Supp Table 8)**. Secondary antibodies were diluted in 1X PBS and applied for 1.5 h at room temperature. Cells were washed with 1X PBS three times prior to incubation with DAPI (1:1,000) for 5 minutes and mounted with Prolong Gold antifade (Thermo Fisher Scientific). All confocal images were captured with a Zeiss LSM 700 laser scanning microscope with the ZEN microscope software. A 63X oil immersion objective with the pinhole set to 1 airy unit was utilized, as well as the same laser power, gain, and zoom for all sets of images and Z-stacks obtained. Z-stacks were obtained with 1024×1024 pixels within a 16-bit range. Image J was used to make representative Z-stack montages and merged Z-stack images^56^. General 20X and 40X immunofluorescence imaging for quantification were performed using an EVOS FL AUTO 2 microscope and the associated-imaging system software.

### Growth and infection of iPSC-derived i^3^Neurons

i^3^Neuron stem cells were maintained at 37 °C in a complete E8 medium (Gibco) on plates coated with Matrigel (Corning) diluted 1:50 in DMEM. Initial three-day differentiation was induced with DMEM/F-12, HEPES (Gibco) supplemented with 1X N2 supplement (Thermo Fisher Scientific), 1X NEAA, 1X Glutamax and 2 μg/ml doxycycline. 10 μM rock inhibitor (Y-27632, Tocris) was added during initial plating of cells to be differentiated and cells were plated onto Matrigel-coated plates. Differentiation medium was replaced daily and after three days of differentiation, partially differentiated neurons were re-plated into cortical neuron (CN) media in coverslips in 24-wells coated with 100 μg/ml poly-L-ornithine (PLO, Sigma). Cortical neuron media consisted of: neurobasal plus medium (Gibco) supplemented with 1X B27 supplement (Gibco), 10 ng/ml BDNF (Peprotech), 10 ng/ml NT-3 (Peptrotech), 1 μg/ml laminin (Gibco) and 1 μg/ml doxycycline. After initial replating, neurons were then maintained in CN media without doxycycline for a further 11 days until mature. Fully differentiated neurons were infected with ZIKV viruses at an MOI 10 in CN media. After 2 hours, the virus inoculum was removed and replaced with 50% fresh – 50% conditioned media. After 96 h p.i., infected cells were fixed 1:1 using 20 mM PIPES pH 6.8, 300 mM NaCl, 10 mM EGTA, 10 mM glucose, 10 mM MgCl_2_, 2% (w/v) sucrose and 5% (v/v) PFA for 10 minutes; and permeabilised for 15 min with 0.3% Triton X-100. Cells were blocked, incubated and imaged as described above.

### Generation, sampling, preparation and infection of human cortical organoids

ALI-COs were prepared as previously described^32,33^. At 82 days *in vitro*, ALI-CO slices were infected with ZIKV at MOI 5 in brain organoid slice media, consisting of: neurobasal medium (Thermo Fisher Scientific) supplemented with 1x B27 supplement (Thermo Fisher Scientific), 0.45% (w/v) glucose (Sigma-Aldrich), 1x Glutamax (Thermo Fisher Scientific) and 1% antibiotic-antimycotic (Thermo Fisher Scientific). After 24 hours, the virus inoculum was removed and replaced with fresh media. Culture medium was replaced by fresh medium every 24 h, and at 7 days post-infection, ALI-COs were fixed with 4% PFA for 3 h. Before cryo-sectioning, ALI-CO tissues were sequentially incubated overnight in 5% and 20% sucrose for cryoprotection and later embedded in OCT. ALI-COs were sectioned at 6 μm thickness. All staining steps were done in permeabilization/blocking buffer (5% BSA, 0.3% Triton X-100 in PBS) at room temperature and their duration was as follows: permeabilization/blocking (1 h); primary antibodies incubation (overnight); wash steps (x3) 10 min each with just PBS 1X; secondary antibodies incubation (1 h); wash steps (x3) 10 min each with just PBS 1X; and DAPI-counterstaining (1:1,000) for 5 min. Sections mounted and cured overnight with Pro-long Gold antifade (Thermo Fisher Scientific) and imaged as described above.

### Mosquitoes

All *in vivo* mosquito experiments used the 15^th^ and 16^th^ laboratory generations of an *Aedes* (*Ae.*) *aegypti* colony originally established from wild specimens caught in Barranquilla, Colombia, in 2017. Mosquitoes were maintained under controlled insectary conditions (28° ± 1 °C, 12 h:12 h light: dark cycle and 70% relative humidity) as previously described^34^. Larvae were reared in dechlorinated tap water supplemented with a standard diet of TetraMin fish food (Tetra). Adults were kept in insect cages (BugDorm) with permanent access to a 10% sucrose solution.

### Mosquito exposure to ZIKV

Mosquitoes were orally challenged with ZIKV by membrane feeding as previously described^34,35^. Briefly, 7-day-old female mosquitoes were starved for 24 h prior to the infectious blood meal. The artificial blood meal consisted of a 2:1 mix of washed rabbit erythrocytes (BCL) and ZIKV suspension containing 6.0 x 10^6^ PFU/mL, supplemented with 10 mM adenosine triphosphate (Merck). The infectious blood meal was offered to mosquitoes for 15 minutes via a membrane-feeding apparatus (Hemotek Ltd.) with porcine intestine as the membrane. Mosquitoes were allowed blood feed for 15 minutes, and fully engorged females were sorted on ice, transferred into 1-pint cardboard containers and maintained under controlled conditions (28° ± 1 °C, 12 h:12 h light: dark cycle and 70% relative humidity) in a climatic chamber with permanent access to 10% sucrose solution for 21 days in the first experiment and 17 days in the second and third experiments.

At 7, 14 and 21 days post-blood feeding in the first experiment and 7, 10, 14 and 17 days post-blood feeding in the second and third experiments, mosquitoes were paralyzed using triethylamine (Sigma) for 5 minutes to collect saliva *in vitro* as previously described^35^. Briefly, the legs of each mosquito were removed, and the proboscis was inserted into a 20-μL pipet tip containing 10 μL of foetal bovine serum (FBS). After 30 minutes of salivation, the saliva-containing FBS was collected and mixed with 40 μL of DMEM (Sigma) containing 2% PenStrep (Gibco) and then stored at −80 °C until use. After the salivation, the head and body remainder of each mosquito were separated with forceps and scalpels and stored individually in separate tubes at −80 °C.

### ZIKV detection in mosquitoes

To assess transmission potential, infectious ZIKV was detected in saliva samples by focus-forming assay in Vero E6 cells as previously described^34^. Briefly, the saliva samples were inoculated onto Vero E6 cells in 96-well plates after removing the cell culture supernatant and incubated at 37 ℃ for 1 hour. The inoculum was removed and replaced by DMEM containing 1.6% carboxymethylcellulose, 1% FBS, 4% Anti-Anti 100X (Gibco) and 1% PenStrep (Gibco). After 5 days of incubation at 37 ℃, cells were fixed with 4% formaldehyde solution (Sigma) for 30 minutes. After fixation, cells were permeabilized with 1X PBS (Gibco) containing 0.1% Triton X-100 (Sigma) for 10 minutes, blocked with 1% BSA (Sigma) in PBS, and then reacted with mouse anti-flavivirus group antigen monoclonal antibody clone D1-4G2-4-15 (Merck) in PBS for 1 h at room temperature (20-25°C). After washing with PBS three times, cells were incubated with a 1:1,000 dilution of Alexa Fluor 488-conjugated goat anti-mouse IgG (Invitrogen) in PBS for 1 h at room temperature. Immunopositive signals were confirmed under a fluorescence microscope EVOS FL (Thermo Fisher Scientific) with appropriate barrier and excitation filters. Transmission potential was determined qualitatively (i.e., positive or negative).

To determine the rates of ZIKV infection and systemic dissemination, mosquito heads and bodies were placed individually in 300 μL of squash buffer containing 10 mM of Tris pH8.0 (Thermo Fisher Scientific), 50 mM of NaCl (Thermo Fisher Scientific) and 1.27 mM of EDTA pH8.0 (Thermo Fisher Scientific) supplemented with proteinase K (Eurobio Scientific) at a final concentration of 0.35 mg/mL and homogenized. A 100-uL sample of the homogenate was transferred to a PCR plate and incubated for 5 minutes at 56 °C, followed by 10 minutes at 98 °C to extract total RNA. Detection of ZIKV RNA was performed by two-step RT-PCR. According to the manufacturer’s protocol, the cDNA was synthesized using M-MLV reverse transcriptase (Thermo Fisher Scientific) with the extracted RNA and random hexameric primers. The synthesized cDNA was subsequently amplified by PCR using DreamTaq DNA polymerase (Thermo Fisher Scientific) with the following primer pair: ZIKV-PCR-Fw (5′-GTATGGAATGGAGATAAGGCCCA-3′) and ZIKV-PCR-Rv (5′-ACCAGCACTGCCATTGATGTGC-3′), according to the manufacturer’s protocol. The cycling conditions were as follows: 2 minutes at 95 °C, 35 cycles of 30 seconds at 95 °C, 30 seconds at 55 °C, and 90 seconds at 72 °C with a final extension step of 7 minutes at 72 °C. Amplicons were visualized by electrophoresis on a 2% agarose gel.

The 5′-UTR mutations of the ZIKV strains were verified by Sanger sequencing in 14-15 head samples for each strain in mosquitoes collected on day 14 post-blood feeding of the second experiment. The synthesized cDNA was amplified using PrimeStar GXL (Takara Bio) with the following primer pair: ZIKV-uORF-PCR-Fw (5′-GTATGGAATGGAGATAAGGCCCA-3′) and ZIKV-uORF-PCR-Rv (5′-TATTGATGAGACCCAGTGATGGC-3′). The cycling conditions were as follows: 2 minutes at 98 °C, 30 cycles of 10 seconds at 98 °C, 15 seconds at 55 °C, and 60 seconds at 68 °C with a final extension step of 2 minutes at 68 °C. Amplified products were directly sequenced in both directions with the following primer pair: ZIKV-uORF-PCR-Fw (same as above) and ZIKV-uORF-SEQ-Rv (5′-GACCCAGCAGAAGTCCGGCTGGC-3′). The expected 5′-UTR sequence was confirmed in all the tested mosquitoes.

### Statistical analysis of results

Data were analysed in GraphPad Prism 9.0 (GraphPad Software, San Diego, CA, USA). Values represent mean ± standard deviation. Statistical significance was evaluated using two-tailed *t*-tests on luciferase data or log_10_(virus titre) data and one-way ANOVA to calculate the percentage of infected cells in brain organoids. Both methods did not assume equal variances for the two populations being compared to calculate the *p*-values. Differences as compared to the control with *p-value* ≤ 0.05 were considered statistically significant, with \**p* < 0.05, ** *p* < 0.01, *** *p* < 0.001 and **** *p* < 0.0001.

The prevalence of ZIKV infection, dissemination and transmission in mosquitoes were analyzed by logistic regression as a function of the experiment, ZIKV strain and time point. The initial statistical model included all their interactions, which were removed from the final model if their effect was non-significant (*p*<0.05). Time was considered a continuous variable. To account for minor differences in the measured virus concentration, the log_10_-transformed blood meal titer was also included in the model as a covariate.

### Ribosomal profiling and RNA-Seq data

10^7^ Vero, U251 or C6/36 cells were grown on 10-cm dishes and infected with parental or mutant viruses at MOI 3. At 24 h p.i., cells treated with CHX (Sigma-Aldrich, 100 μg/ml) were incubated for 3 minutes and then rinsed with 5 ml of ice-cold PBS before being flash-frozen. Cells harvested by ‘flash-freezing’ were directly rinsed with 5 ml of ice-cold PBS, flash-frozen in a dry ice/ethanol bath, and lysed with 400 μl of lysis buffer (20 mM Tris-HCl pH 7.5, 150 mM NaCl, 5 mM MgCl_2_, 1 mM DTT, 1% Triton X-100, 100 μg/ml cycloheximide and 25 U/ml TURBO DNase). The cells were scraped extensively to ensure lysis, collected and triturated ten times with a 26-G needle. Cell lysates were clarified by centrifugation at 13,000 *g* for 20 min at 4 °C. Lysates were subjected to Ribo-Seq and RNA-Seq based on previously reported protocols^17^. Vero and U251 ribosomal RNA was removed using Ribo-Zero Gold rRNA removal kit (Illumina); and C6/36 ribosomal RNA contamination was removed by treatment with duplex-specific nuclease (DSN) as previously described^57^. Library amplicons were constructed using a small RNA cloning strategy adapted to Illumina smallRNA v2 to allow multiplexing. Amplicon libraries were deep-sequenced using an Illumina NextSeq500 platform. Ribo-Seq and RNA-Seq sequencing data have been deposited in the ArrayExpress database (http://www.ebi.ac.uk/arrayexpress).

### Computational analyses of sequence data

Reads were demultiplexed. Adaptors were trimmed off using the FASTX-Toolkit version 0.0.14 (http://hannonlab.cshl.edu/fastx_toolkit/), and sequences shorter than 25 nt after trimming were discarded. Sequences not linked to an adaptor and sequences constituted of adaptors ligated together were also removed. Reads remaining after this initial selection were sequentially mapped to (i) ribosomal RNA (rRNA); (ii) the American or African ZIKV genomic RNA (viral genome sequences were confirmed by *de novo* assembly using Trinity version2.8.5); (iii) messenger mRNA (mRNA); (iv) non-coding RNA (ncRNA); (v) genomic DNA (gDNA) and (vi) a contaminants database, using bowtie (version 1.2.3) with parameters -v n_mismatches --best (i.e., maximum of n_mismatches mismatches permitted, report best match) where n_mismatches was 1 for mapping in U251 cells and 2 for Vero and C6/36 cells. The remaining reads were classified as unmapped. The rRNA databases included the Genbank accession numbers listed in **Supp Table 9**. The mRNA sequences for each host were downloaded from NCBI RefSeq. The ncRNA databases for *C. sabaeus* were retrieved from Ensembl release 91 and compiled with 534 tRNA sequences of *M. mulatta* and Genbank accession XM_007980346; for *H. sapiens,* the ncRNA sequences were downloaded from Ensembl release 105 and for *A. albopictus* from Ensembl release 55. The gDNA sequences for each host were retrieved from the same Ensembl releases as the ncRNA. The contaminants database comprises Genbank accessions of potential contaminants that might be found in the lab and are used as quality control to ensure that no/very few reads map to this database. The database includes viruses (e.g., HSV-1, PRRSV) and bacteria (e.g., *E. coli*) that are regularly used within the Division of Virology, University of Cambridge, and other common potential contaminants (e.g., mycoplasma sequences). The composition of libraries (**Supp Fig. 9**) indicated some contamination of Vero Dak84 RNA-Seq (replicate 1 and 2) with Toscana virus (TOSV). These reads do not exhibit the same features as the rest of the reads in those libraries (i.e., length distribution of the reads does not have the same shape), indicating that the contamination occurred after lysates were harvested and thus, they do not affect our conclusions.

Quality control analyses of the reads (**Supp Figs. 10-13**), including length distribution, phasing, and phasing per read length, were carried out using Ribo-Seq and RNA-Seq reads mapping to host mRNA and viral RNA as previously described^17^. To assess ribonucleoprotein (RNP) contamination of Ribo-Seq samples, a comparison was made between the number of reads mapping to CDSs and the 3′ UTR of host mRNAs only (**Supp Fig. 14**).

For the visualisation of read densities on the viral genome, the position of the P-site was calculated by adding a fixed +12 nt offset to the 5′ end of each read. Read densities were normalized by the sum of total viral RNA and total host mRNA and plotted in reads per million mapped reads (RPM). The genomic regions with coordinates 15-125 or 18-315 for the parental and mutant viruses, respectively, were chosen for the zoom plots of the 5′ UTR. The region from genomic coordinate 10,370 to the end of the genome was used for the zoom plots of the 3′ UTR. Summarising bar charts show the percentage of reads in each phase, normalised by the total number of reads in the designated region (**Supp Table 10**). The reads per kilobase per million mapped reads (RPKM) values for Ribo-Seq were divided by the corresponding RNA-Seq values to calculate the translational efficiency (TE).

## Acknowledgements

The authors are indebted to Andrew Firth (Department of Pathology, University of Cambridge) for all the intellectual input and technical support in the bioinformatic analysis, and for the critical reading of this manuscript. The authors would like to thank Alain Kohl (Centre for Virus Research, University of Glasgow) and Lindomar J. Pena and Rafael Oliveira de Freitas França (Fiocruz Recife, Pernambuco, Brazil) for the provision of PE243 ZIKV RNA used to generate the virus stock; and Andres Merits (University of Tartu, Tartu, Estonia) for providing the pCC1-SP6-Am ZIKV plasmid. The African ZIKV isolate (Dak84) was provided by the ‘European Virus Archive goes Global (EVAg) project’ funded by the European Union’s Horizon 2020 research and innovation programme under grant agreement No 653316. We thank Catherine Lallemand for assistance with mosquito rearing. We are grateful to Claudia Romero-Vivas, who initially provided the mosquito colony from Colombia. NI would like to thank David Carpentier (Confocal microscopy facility, Department of Pathology, University of Cambridge) and Louise Howard (Histology Facility, Department of Pathology, University of Cambridge) for their helpful assistance.

C.L. was supported by the ‘Wolfson-Pathology PhD studentship’ (University of Cambridge). G.M.C. was supported by a Wellcome Trust grant [203864/Z/16/Z]. A.M.D. and H.S. were supported by the Wellcome Trust Senior Research Fellowships [106207/Z/14/Z, 220814/Z/20/Z] and a European Research Council grant [646891] awarded to Andrew Firth (Department of Pathology, University of Cambridge). J.E.D. and A.S.N. were supported by a Wellcome Trust Senior Research Fellowship [219447/Z/19/Z] awarded to J.E.D. L.W.M. and I.G. were supported by a Wellcome Trust grant [207498/Z/17/Z] to I.G. Sir Henry Dale Fellowship [220620/Z/20/Z] from the Wellcome Trust and the Royal Society, MRC project grant [MR/T000376/1] and an Isaac Newton Trust/Wellcome Trust ISSF/University of Cambridge Joint Research Grant to V.L. Research in the A.L. laboratory was funded by the Medical Research Council UK (UKRI: MR/P008658/1; MR/X006867/1 awarded to A.L.). This work was also supported by the French Government’s Investissement d’Avenir program Laboratoire d’Excellence Integrative Biology of Emerging Infectious Diseases (grant ANR-10-LABX-62-IBEID to L.L.) and MSDAVENIR (grant INTRANZIGEANT to L.L.). S.T. was supported by an MSCA fellowship from the European Union (grant MSCA-2021-PF-01 ZIKVMosTransmit). I.B. was supported by grants from the Wellcome Trust [202797/Z/16/Z] and the Biotechnology and Biological Sciences Research Council UK [BB/V000306/1]. N.I. was supported by an Isaac Newton Trust Grant [18.40r], a Royal Society Research Grant [RGS\R1\191137], an Isaac Newton Trust/Wellcome Trust ISSF/University of Cambridge Joint Research Grant and partially by the NIH Grant [5R21AI147172].

