## Supplementary Information for "Zika viruses encode multiple upstream open reading frames in the 5′ viral region with a role in neurotropism"

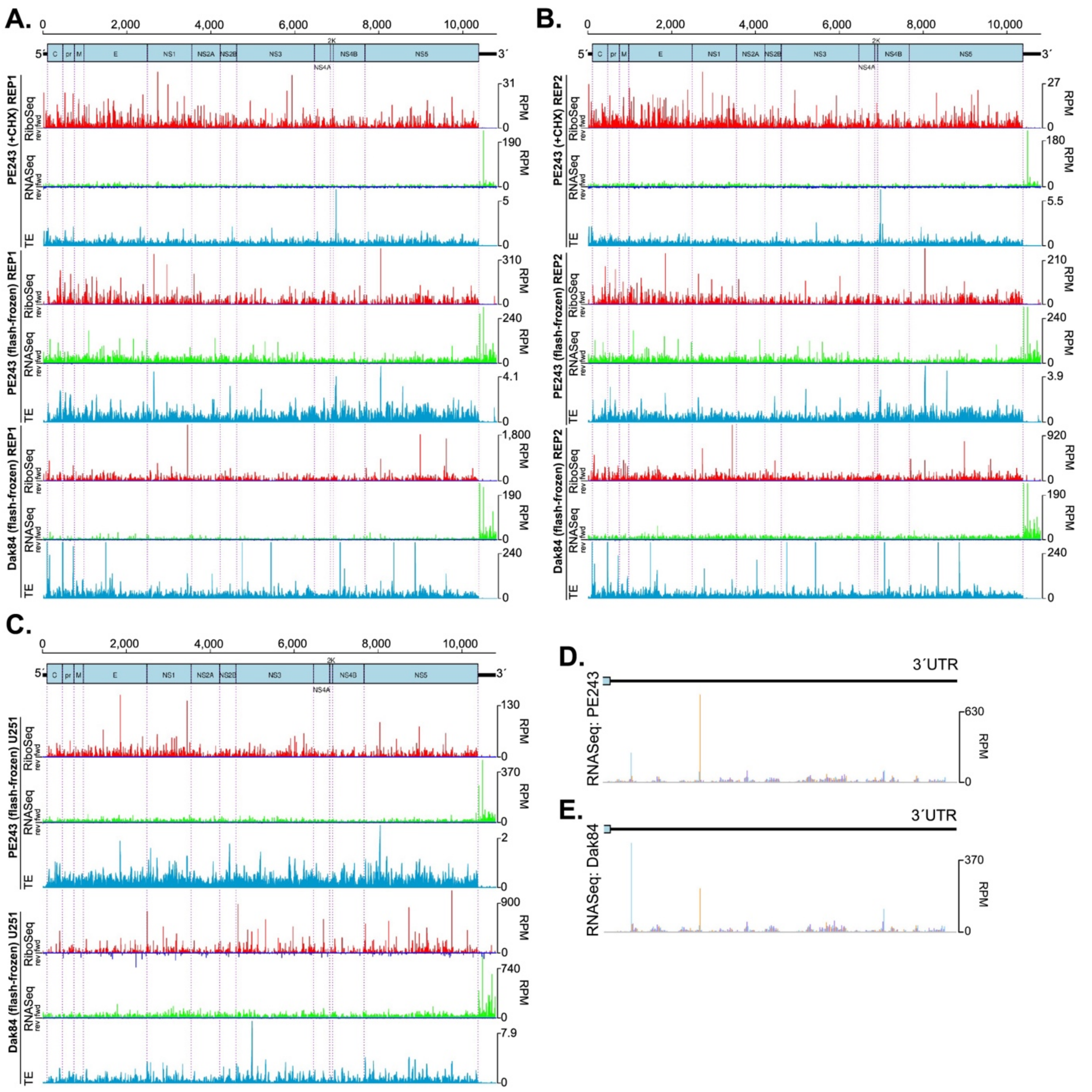

**Supp. Figure 1. ZIKV RNA synthesis and translation.** (**A**) Ribo-Seq (red) and RNA-Seq (green) densities in reads per million mapped reads (RPM) of repeat 1 (REP1) of PE243 (MOI:3) at 24 h p.i. in Vero cells pre-treated with CHX (upper panel), flash-frozen Vero cells (middle panel); or flash-frozen Vero cells infected with Dak84 (MOI:3) at 24 h p.i. (lower panel) as described in **Fig 1**. TE (light blue) is translational efficiency. Vero cells pre-treated with CHX panel is a duplication of **Fig 1B** for coherence. (**B**) Repeat 2 (REP2) as in **A**. (**C**) Flash-frozen U251 cells infected with PE243 (MOI:3, upper panel) or Dak84 (MOI:3, lower panel) at 24 h p.i. (**D-E**) Zoom plot of RNA-Seq densities at 24 h p.i. for Vero cells in the PE243 (**D**) or Dak84 (**E**) 3′ UTR.

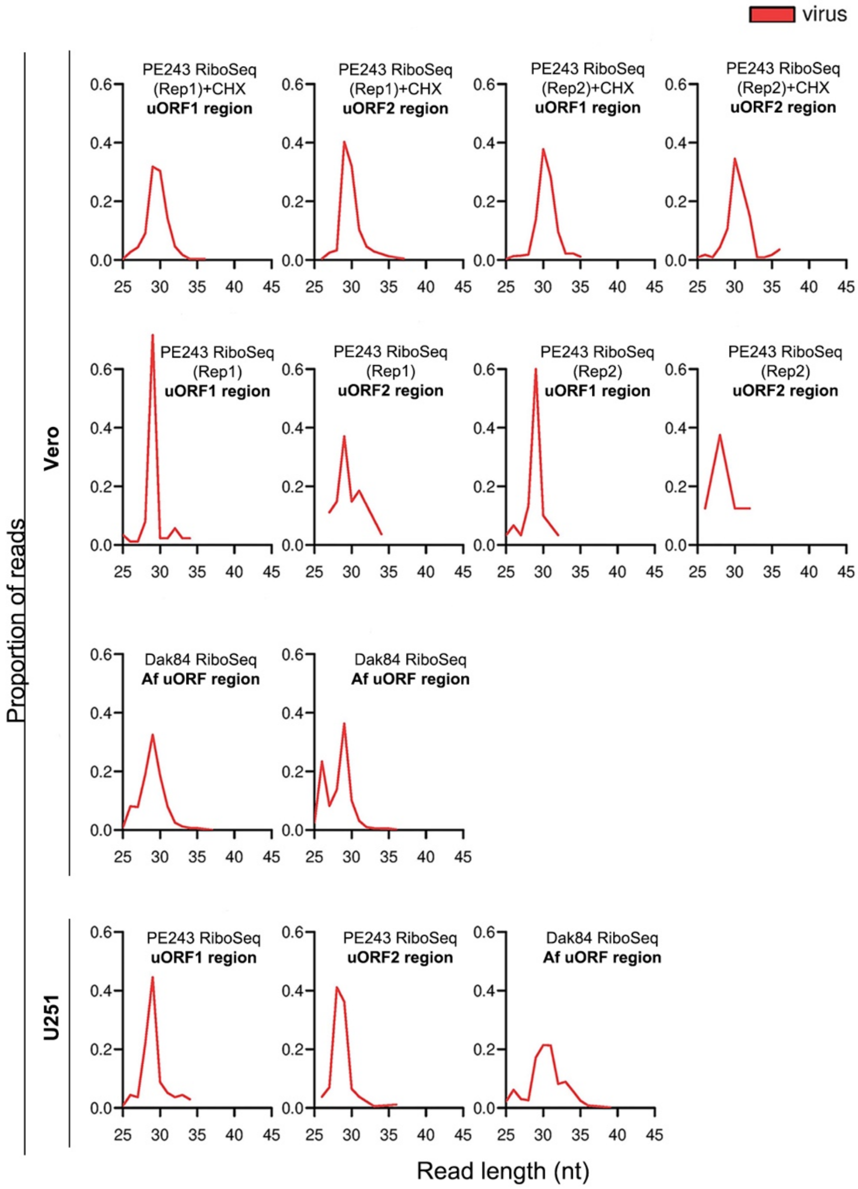

**Supp. Figure 2. Length distribution and proportion of Ribo-Seq reads mapping to the different uORFs** (in red) for Vero and U251 cells infected with PE243 and Dak84.

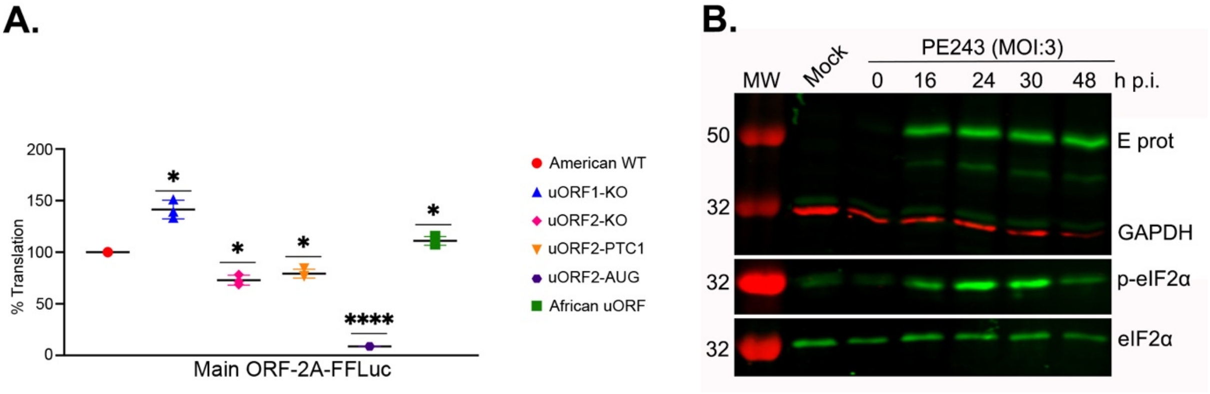

**Supp. Figure 3. Analysis of ZIKV uORF translation using luciferase reporters.** (**A**) Relative FF-Luc/Ren-Luc activity of main-ORF-2A-FFLuc mutant reporters in U251 cells. Cells were harvested at 30 h p.t.. Experiments were performed in triplicate with three biological replicates. All *t*-tests were two-tailed and did not assume equal variance for the two populations being compared (**p* < 0.05, **** *p* < 0.0001). All *p*-values are from comparisons of the mutant with the wild-type. (**B**) Western blot analysis of Vero cells infected with PE243 (MOI:3) and harvested at 0, 16, 24, 30 and 48 h p.i. Cell lysates were probed against viral E protein (E prot) and the phosphorylated version of eIF2α. GAPDH and eIF2α are used as loading controls. Molecular masses (in kDa) are indicated on the left.

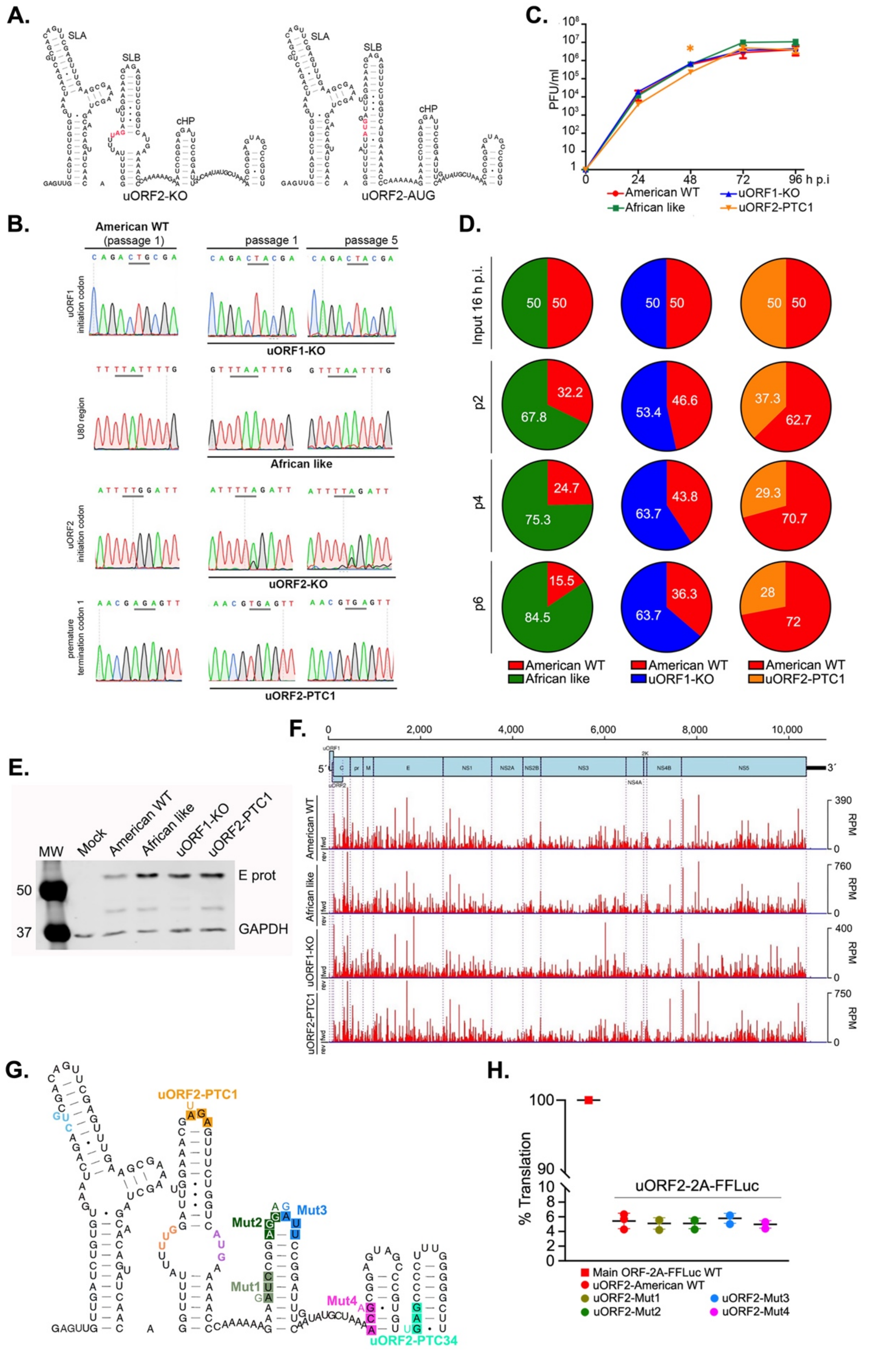

**Supp. Figure 4. The significance of uORFs translation in virus infection.** (**A**) SHAPE RNA secondary structure of the 5′ region (first 180 nucleotides) of the uORF2-KO and uORF2-AUG mutant viruses. Modified codons are indicated in red. SLA (stem-loop A), SLB (stem-loop B) and cHP (capsid hairpin). (**B**) Sequencing histograms of RT-PCR products of the American

WT, uORF1-KO, African-like, uORF2-KO and uORF2-PTC1 viruses at passage 1 and passage 5. Mutated nucleotides are underlined. (**C**) Time-course of Vero cells infected with ZIKV mutant viruses (MOI 0.01) for 96 h. Plaque assays were performed as described in **Fig 3C**. All *t*-tests were two-tailed and did not assume equal variance for the two populations being compared ($*p < 0.05$). All *p*-values are from comparisons of the mutant virus with the American WT. (**D**) Pie charts of the competition assays of American WT and mutant viruses at 50:50 proportion in Vero cells as described in **Fig 3D**. Experiments were repeated independently eight times (raw data in **Supp Table 5**). (**E**) Western blot analysis of Vero cells infected with the American WT, African-like, uORF1-KO and uORF2-PTC1 viruses (MOI:3) for 24 h. Cellular extracts were probed against viral E protein and GAPDH (as loading control). Molecular masses (kDa) are indicated on the left. (**F**) Ribo-Seq density, in reads per million mapped reads (RPM), at 24 h p.i. in flash-frozen Vero cells infected with American WT, African-like, uORF1-KO and uORF2-PTC1 viruses (MOI:3) as described in **Fig 1B**. (**G**) Scheme of the 5′ region of the American WT. Premature termination codons (PTC) are colour-squared in orange and bright green for uORF2-PTC1 and uORF2-PTC34, respectively. Nucleotides corresponding to 'alternative' non-canonical initiation codons for uORF2 are colour-coded and named (Mut1-Mut4). Mutated nucleotides are located next to the original. (**H**) Relative FF-Luc activity for the different mutants of uORF2-2A-FFLuc in Vero-transfected cells as in **Fig 2B**. 100% translation accounted for the main ORF WT (red square).

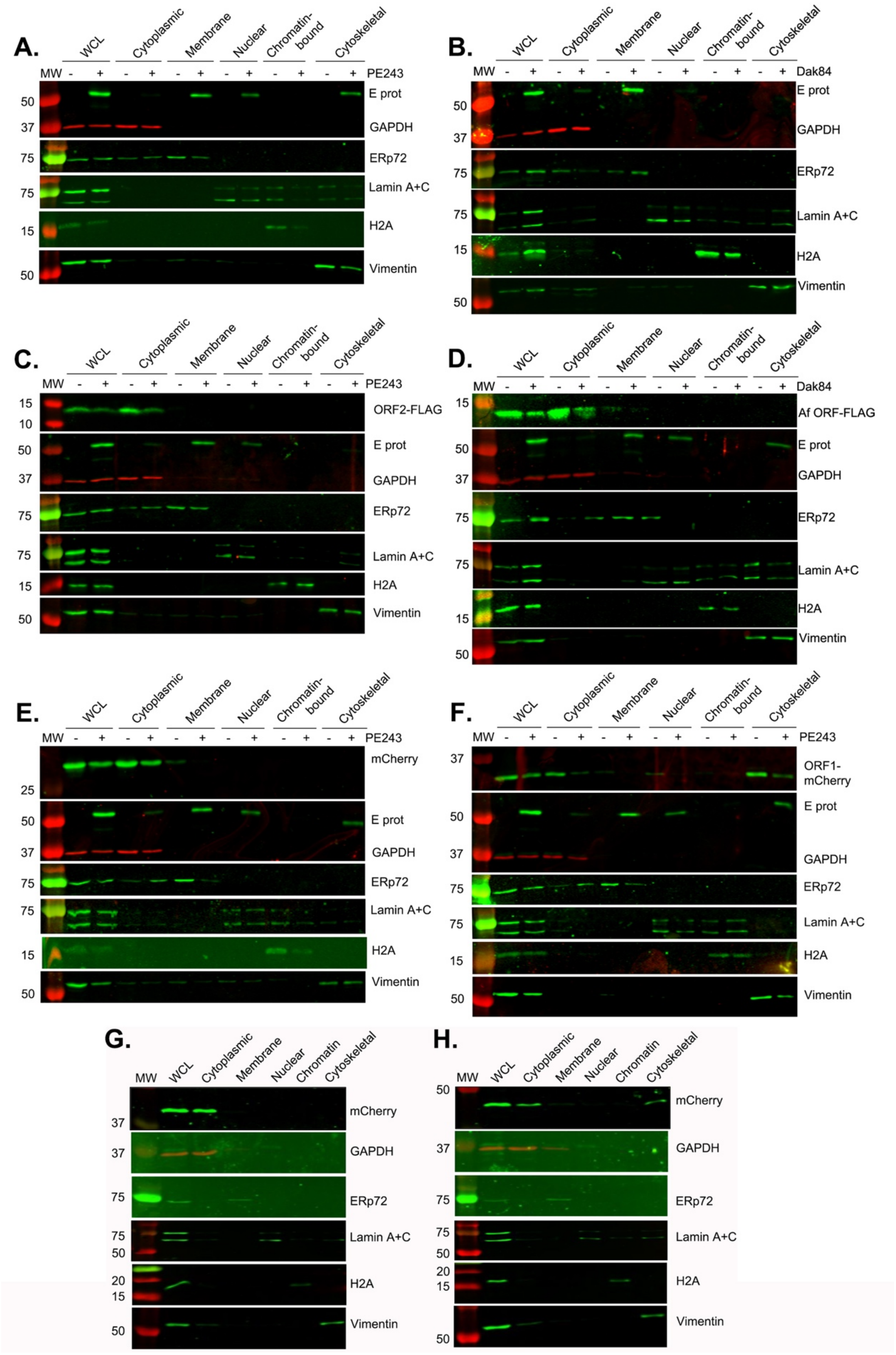

**Supp. Figure 5. Functional characterization of ZIKV uORF-encoded proteins.** (A-F) Vero cells were transfected with the corresponding pCAG plasmid for 24 hours and infected with either PE243 or Dak84 (MOI:1). Cells were harvested at 14 h p.i. (Dak84 infection) or 16 h p.i. (PE243 infection) and subjected to subcellular fractionation. Western blot analysis of total extract (WCL), cytoplasmic, membrane, nuclear, chromatin and cytoskeletal fractions was carried out as follows: membranes were probed with antibodies against FLAG or mCherry for detecting the *in trans* tagged protein; E protein (ZIKV infection); GAPDH (cytosolic marker); ERp72 (membrane marker); Lamin A+C (nuclear marker); H2A (chromatin marker); and vimentin (cytoskeletal marker). Molecular masses (kDa) are indicated on the left. Combinations tested were pCAG-empty plasmid plus PE243 (**A**); pCAG-empty plasmid plus Dak84 (**B**); pCAG-uORF2-FLAG plus PE243 (**C**); pCAG-African ORF-FLAG plus Dak84 (**D**); pCAG-mCherry plus PE243 (**E**) and pCAG-uORF1-mCherry plus PE243 (**F**). U251 cells were transfected with pCAG-mCherry (**G**) and pCAG-uORF1-mCherry (**H**) for 40 h p.t. and subjected to subcellular fractionation and analysis as described above.

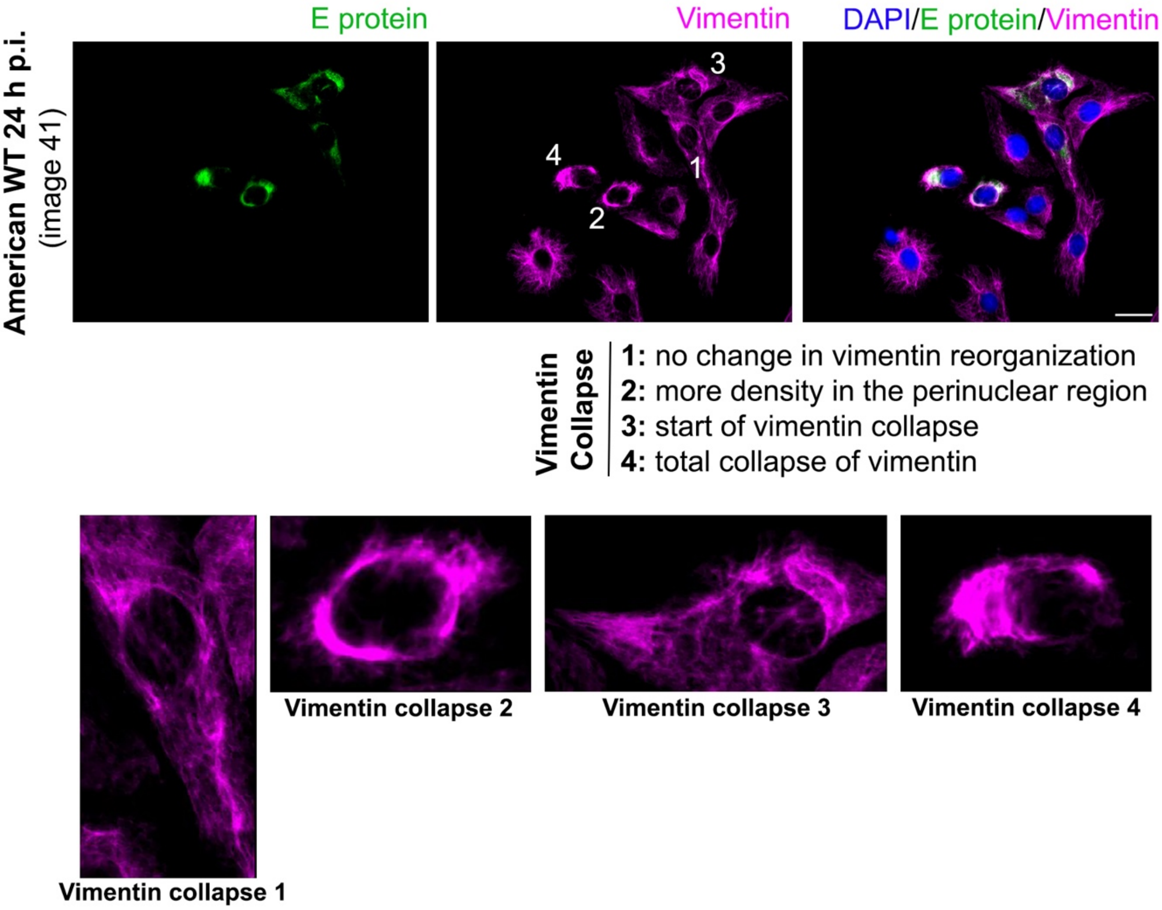

**Supp. Figure 6. The collapse of vimentin by uORF1.** Representative immunofluorescence microscopy image (American WT 24 h p.i., image 41) showing the four stages of vimentin collapse in Vero-infected cells as indicated in the figure. Cells were stained for viral E protein (green), vimentin (magenta) and nuclei (DAPI in blue).

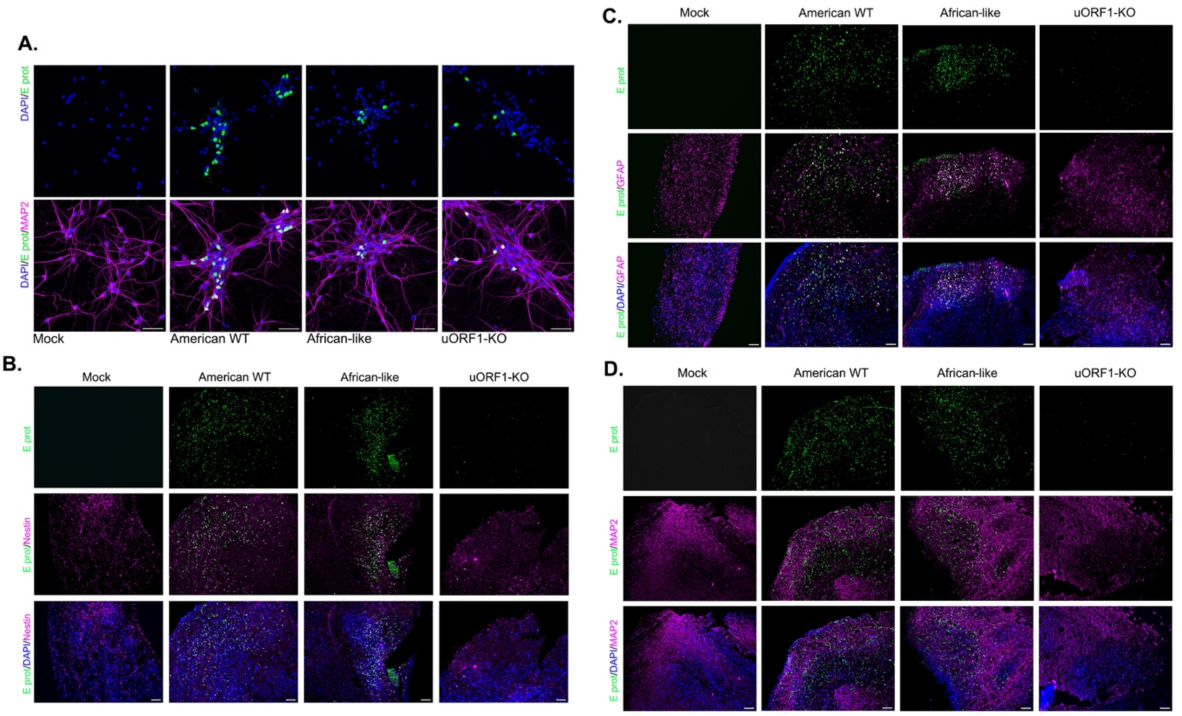

**Supp. Figure 7.** (**A**) Representative confocal images of i$^3$Neurons infected with the American WT, the African-like and the uORF1-KO viruses (MOI:10) for 96 h. Cells were stained with antibodies against the viral E protein (green) and MAP2 (magenta), a marker for cortical neurons. Nuclei were counter-stained with DAPI (blue). Scale bars, 50 µm. (**B**-**D**) Representative images (10X resolution) of ALI-COs infected with the American WT, the African-like and the uORF1-KO viruses (MOI:5) for 7 days showing immunoreactivity for the viral E protein (green) and different cellular markers (magenta), including nestin (**B**), GFAP (**C**) and MAP2 (**D**). Nuclei were counter-stained with DAPI (blue). Scale bars, 100 µm.

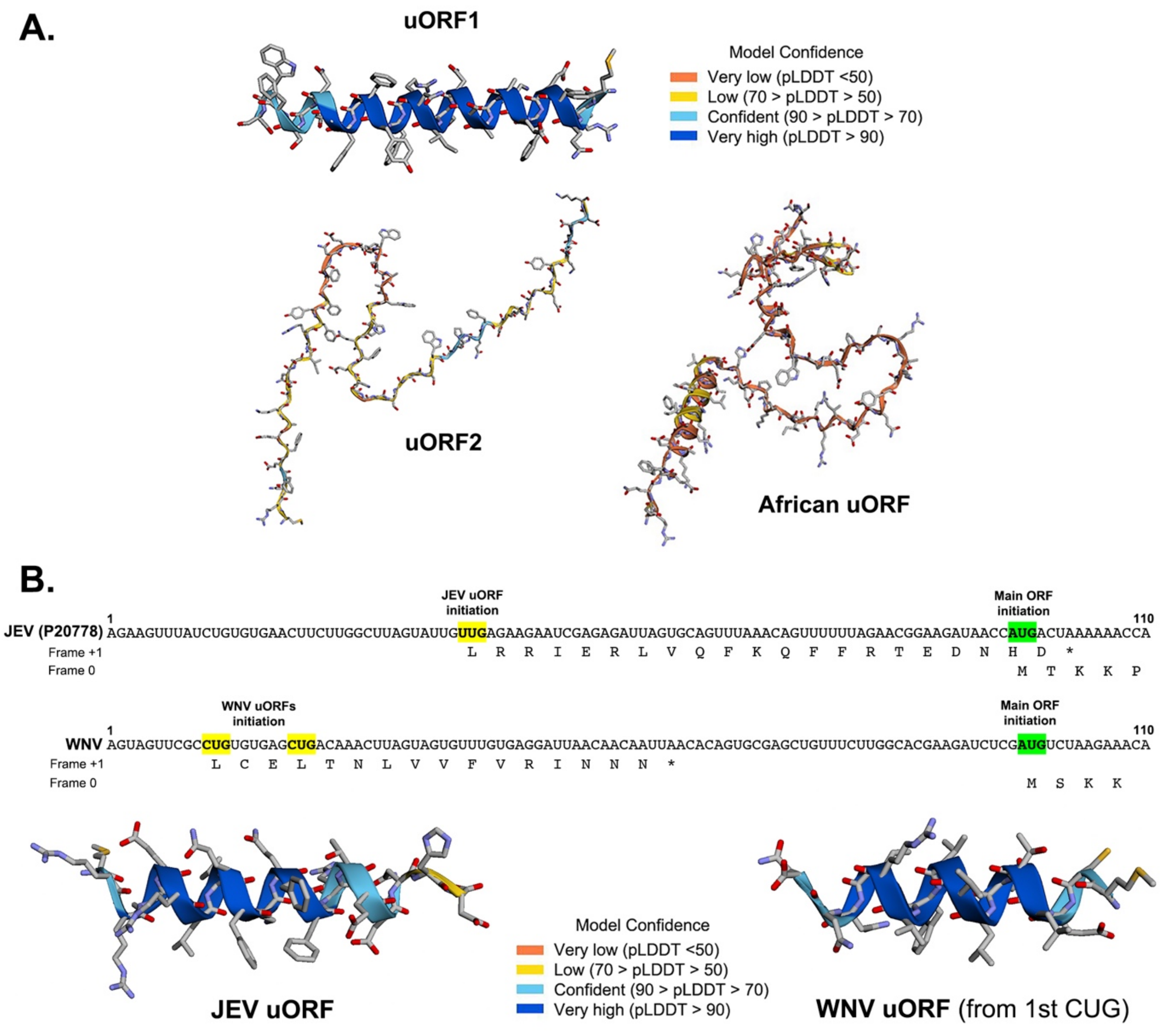

**Supp. Figure 8.** (A) **AlphaFold2 predictions for ZIKV uORF-encoded proteins**. uORF1 is predicted to form an α-helix. uORF2 yields a low confidence prediction and is likely to be intrinsically disordered, but it could become ordered upon binding cellular partners. The African uORF, which comprises the American uORF1 and uORF2, also yields a low confidence prediction consistent with little intrinsic structure, although the N terminus is predicted with low confidence to have an α-helical structure similar to uORF1. (**B**) **AlphaFold2 predictions for JEV uORF- and WNV uORF-encoded proteins.** Ribosome profiling of JEV (P20778 strain) reveals translation of an uORF in the 5′ region using a non-canonical initiation codon (UUG in yellow)[43]. The JEV uORF-encoded peptide (primary sequence indicated in the +1 frame) is predicted to form an α-helix with high confidence as ZIKV uORF1. The 5′ region of WNV (accession number M12294.2) has two non-canonical initiation codons (in yellow) that could code for uORFs longer than 10 residues. The second CUG initiation codon aligns with the ZIKV uORF1 CUG initiation codon. The WNV uORF-encoded peptide initiating from the 1st CUG (primary sequence indicated in the +1 frame) is also predicted with high confidence to form an α-helix. The AUG initiation codon for the main ORF translation is indicated in green.

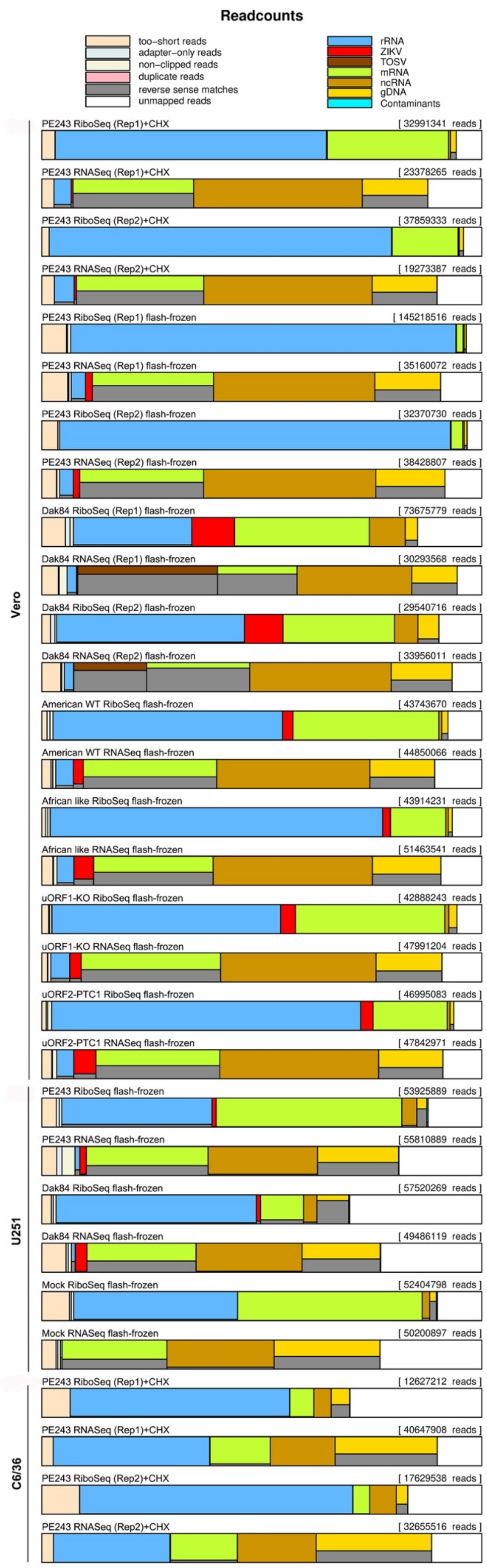

**Supp. Figure 9. Composition of libraries.** Reads were mapped to ZIKV RNA, Toscana virus (TOSV) RNA, host rRNA, mRNA, ncRNA and gDNA databases, and our contaminant database. Reads mapping to gDNA are expected to derive from unannotated transcripts not present in the mRNA or ncRNA databases, but, since the direction of transcription is not annotated in the gDNA database, such reads constitute a mixture of forward and reverse-sense matches. Reverse-sense rRNA matches in the RNA-Seq samples are expected to derive from rRNA depletion kits which contain complementary sequences to rRNA.

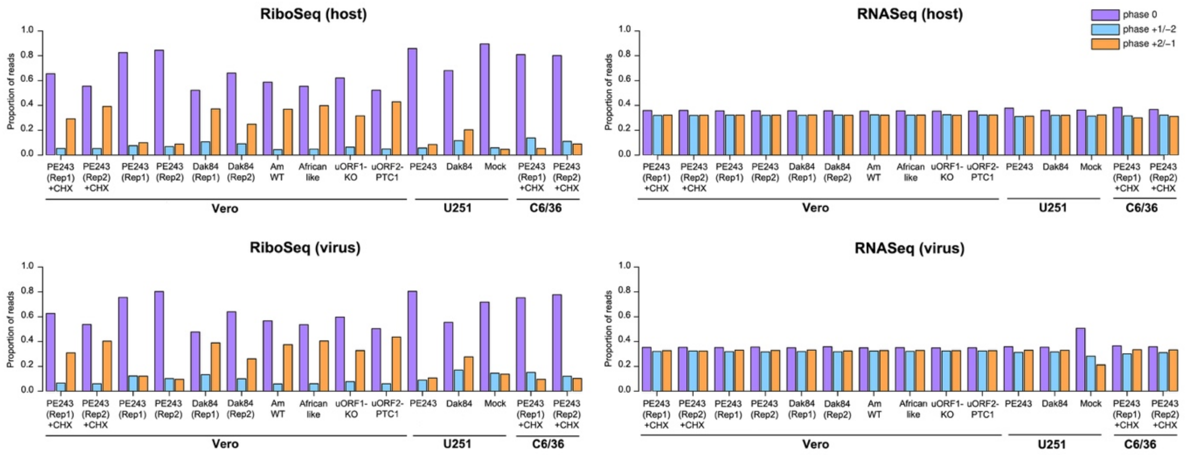

**Supp. Figure 10. Phasing of reads.** Proportion of reads (all read lengths) attributed to each phase, for positive-sense reads mapping within host CDSs (upper) and virus (lower). Phases correspond to which position within the codon the 5′ end of the read maps to (0: purple, 1: blue, 2: orange). The 5′ end coordinate of Ribo-Seq reads is influenced by the position of the translating ribosome, leading to a clear dominance of the 0 phase. For RNA-Seq reads, the 5′ end coordinate is determined by alkaline hydrolysis, so it does not result in a dominant phase. Note that the low virus read count for mock samples in the virus plots (lower) makes the phase distributions prone to noise.

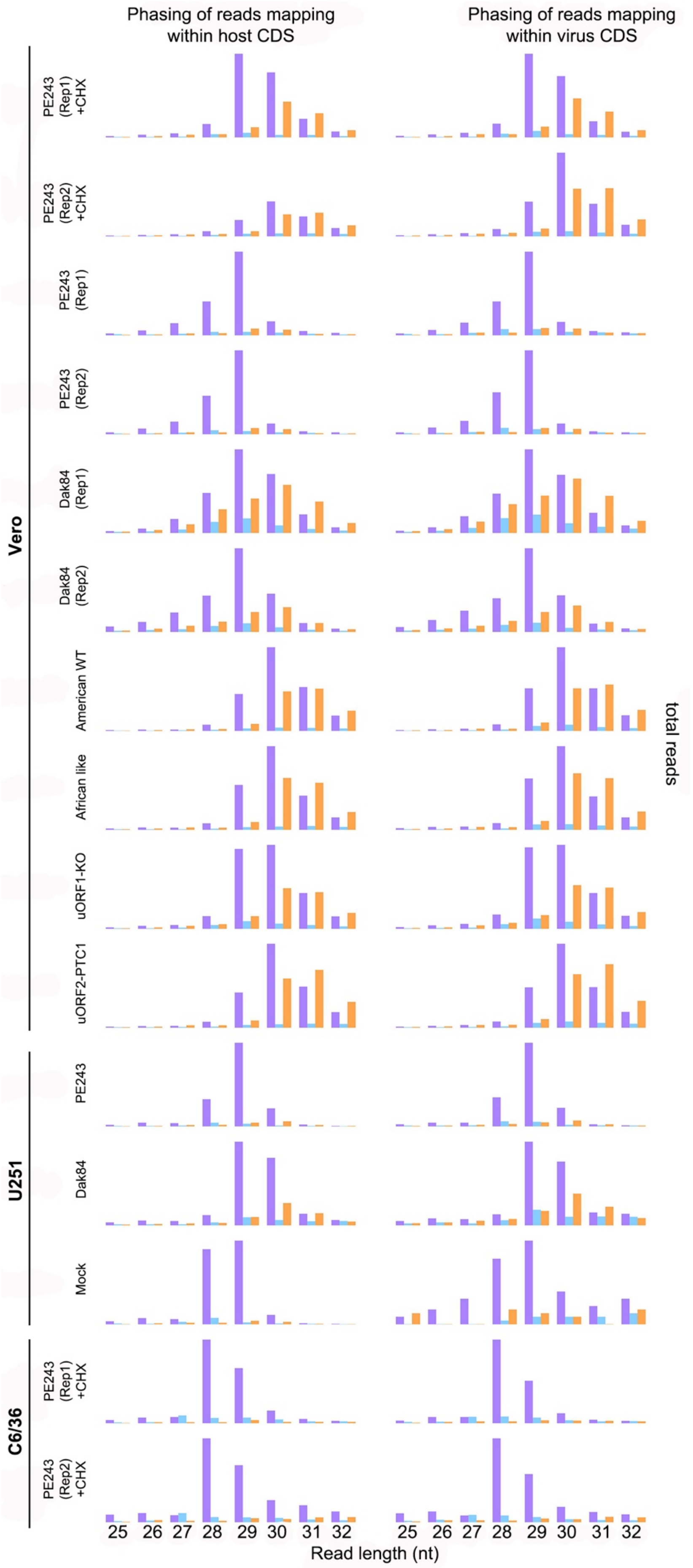

**Supp. Figure 11. Phasing of reads distributed by read length** (from 25 to 32 nt). Phasing shown as in Supp. Figure 10.

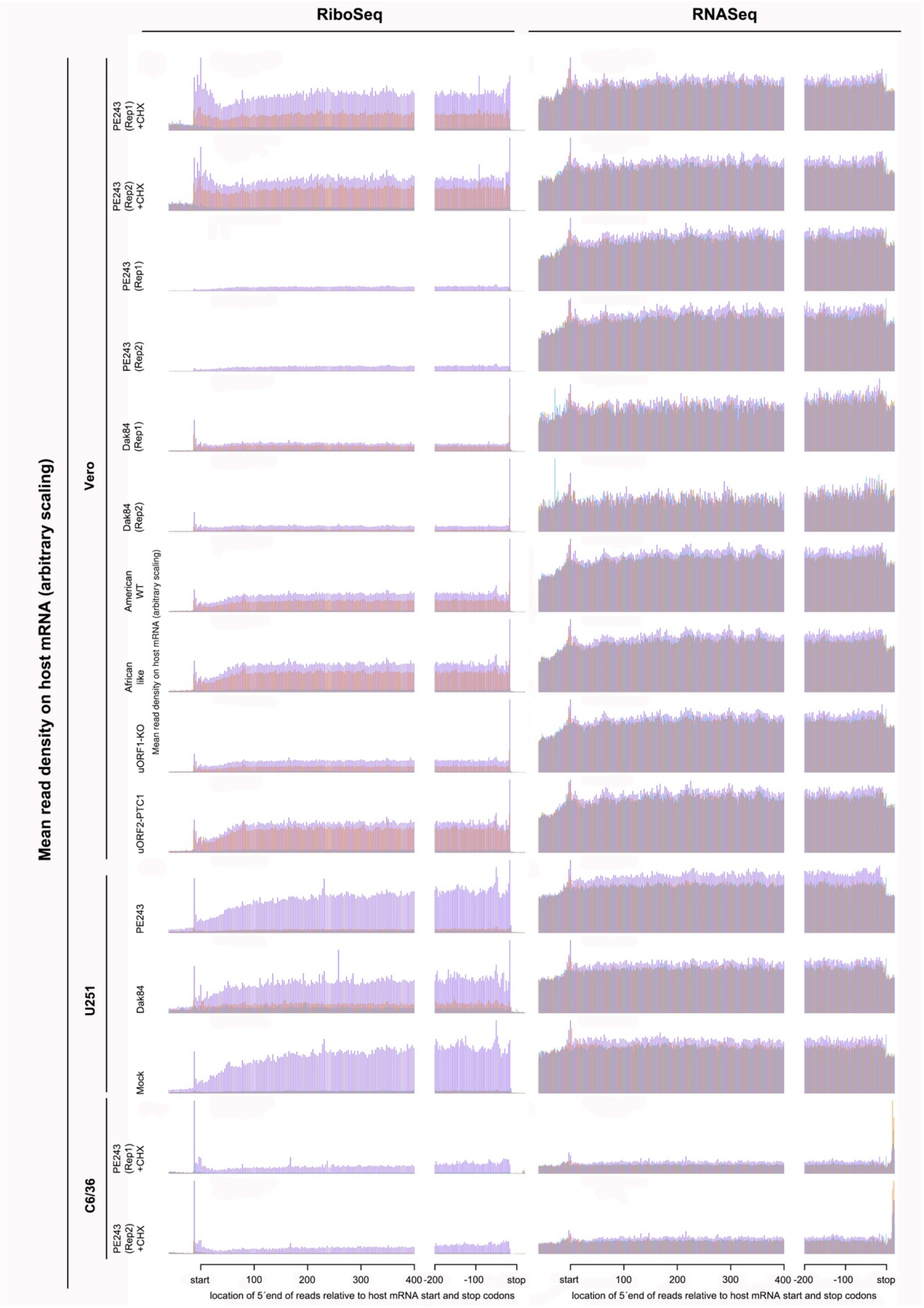

**Supp. Figure 12. RPF and RNA-Seq distributions on host mRNAs.** Histograms of RPF (left) and RNA-Seq read (right) 5′ end positions relative to annotated initiation and termination codons summed over all host RefSeq mRNAs for the Ribo-Seq (left) and RNA-Seq (right) libraries. To account for different library sizes, histograms are normalised by the sum of total virus RNA (positive and negative-sense) plus total host mRNA for the library.

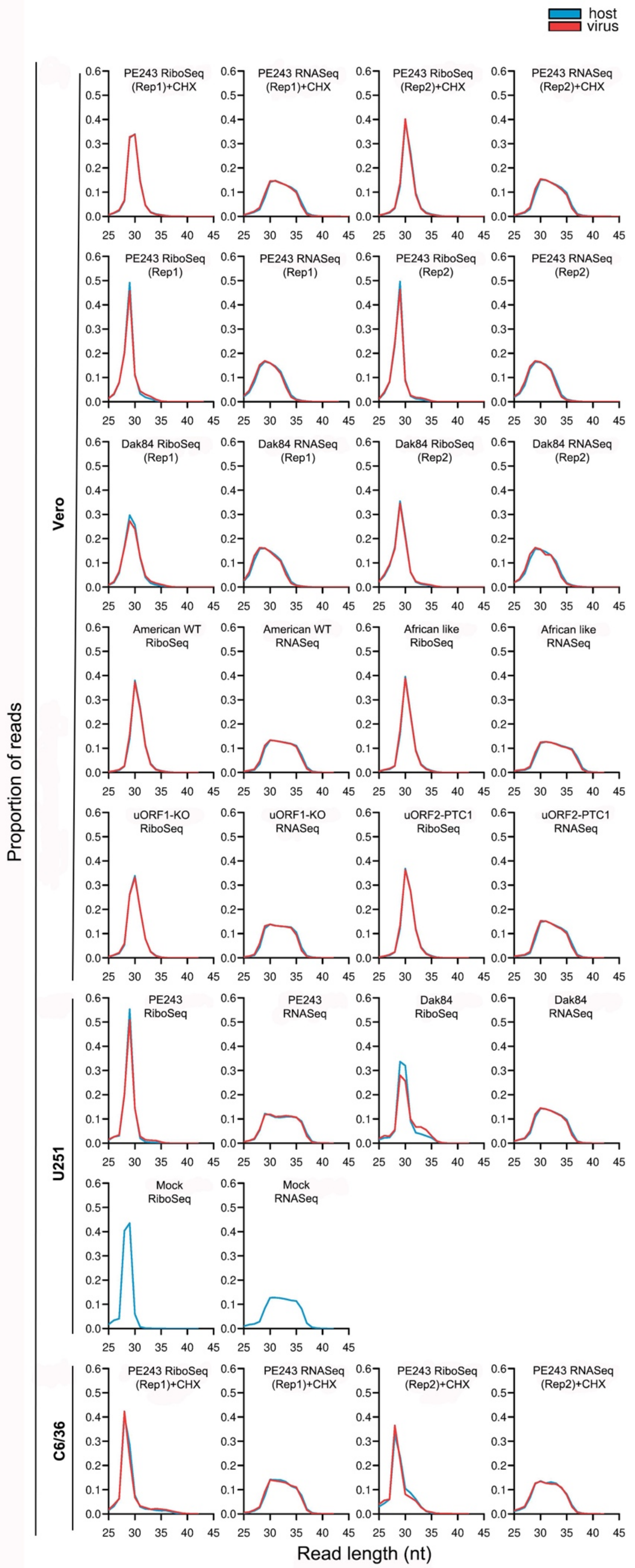

**Supp. Figure 13. Comparison of read length distributions for virus and host mRNA.** Length distributions for reads mapping to positive-sense host mRNAs (blue) and virus RNA (red). Every panel in each pair shows the distributions normalized to have equal total sums to facilitate the comparison of distribution shapes. Differences between host and virus distributions are indicative of contamination.

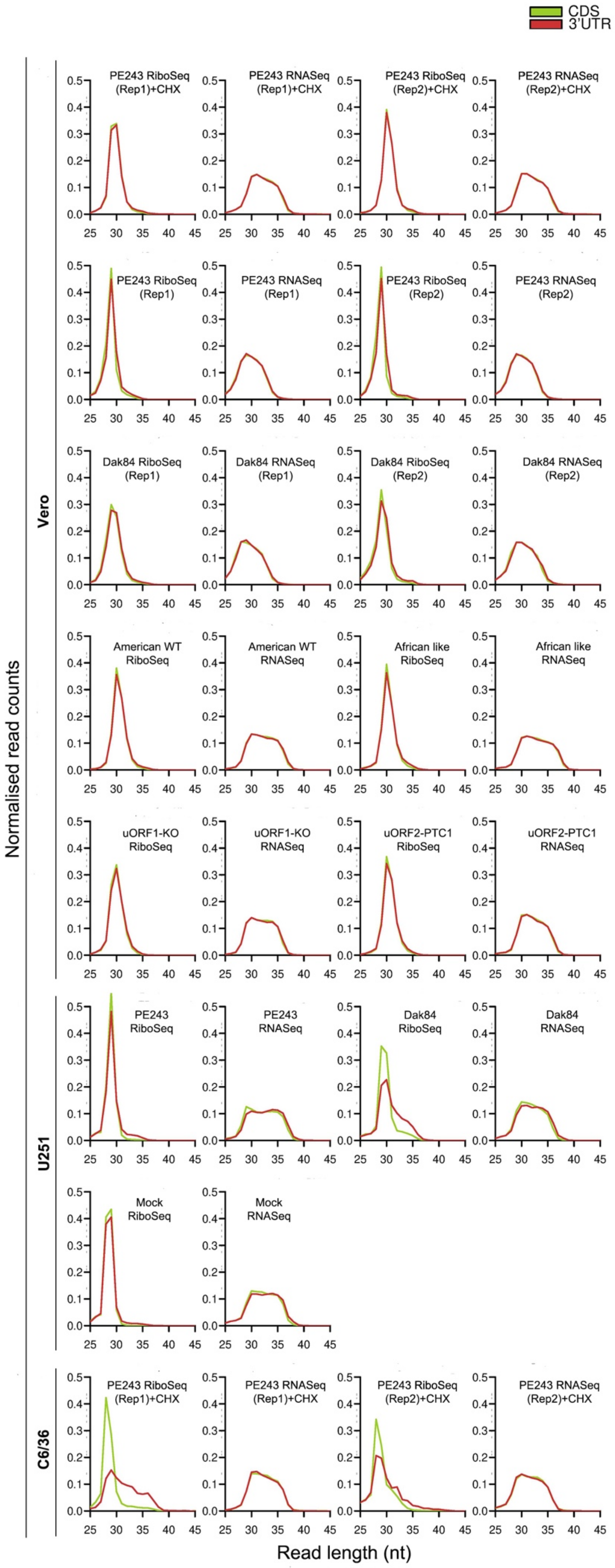

**Supp. Figure 14. Comparison of the length distribution of reads mapping to host mRNA coding regions and 3′ UTRs.** Reads were counted in windows from 10 to 100 codons upstream (CDS; green) or downstream (3′ UTR; red) of annotated termination codons, and summed over all host mRNAs. Every panel in each pair shows the distributions normalized to have equal total sums so that the shapes of the CDS and 3′ UTR distributions can be compared. For RNA-Seq, the two distributions have essentially identical shapes. For Ribo-Seq, differences in the two distributions provide an indicator of the level of non-RPF contamination present in the sample.

### Supplementary Table 1: ZIKV T7 Luciferase reporters: Vero cells

#### Replicate 1

##### FFLuc 24 h p.i.

| | | | |
|---|---|---|---|
| Mock | 20 | 20 | 40 |
| pREN | 80 | 50 | 90 |
| ORF1 FF | 569700 | 479500 | 429300 |
| ORF2 FF | 1753000 | 2047000 | 1402000 |
| Main ORF FF | 36590000 | 34060000 | 32030000 |

##### Ren Luc

| | | | |
|---|---|---|---|
| Mock | 50 | 100 | 50 |
| pREN | 258900 | 244100 | 265900 |
| ORF1 FF | 307700 | 288300 | 283600 |
| ORF2 FF | 222900 | 281200 | 221400 |
| Main ORF FF | 313100 | 329300 | 316900 |

##### FFLuc 24 h p.i. + PE243 MOI 3

| | | | |
|---|---|---|---|
| Mock | 20 | 40 | 30 |
| pREN | 90 | 50 | 50 |
| ORF1 FF (PE243) | 430000 | 449600 | 338900 |
| ORF2 FF (PE243) | 2073000 | 1704000 | 1180000 |
| Main ORF FF (PE243) | 27820000 | 33230000 | 24440000 |

##### Ren Luc

| | | | |
|---|---|---|---|
| Mock | 80 | 60 | 70 |
| pREN | 289900 | 227200 | 317700 |
| ORF1 FF | 166000 | 196400 | 151900 |
| ORF2 FF | 206000 | 212000 | 111900 |
| Main ORF FF | 191700 | 259500 | 181600 |

#### Replicate 2

##### FFLuc 24 h p.i.

| | | | |
|---|---|---|---|
| Mock | 20 | 20 | 40 |
| pREN | 80 | 50 | 90 |
| ORF1 FF | 482400 | 463500 | 240400 |
| ORF2 FF | 1538000 | 1535000 | 728900 |
| Main ORF FF | 24790000 | 27170000 | 16360000 |

##### Ren Luc

| | | | |
|---|---|---|---|
| Mock | 50 | 100 | 50 |
| pREN | 258900 | 244100 | 265900 |
| ORF1 FF | 283000 | 288700 | 194300 |
| ORF2 FF | 233900 | 290100 | 152300 |
| Main ORF FF | 299100 | 309600 | 215600 |

##### FFLuc 24 h p.i. + PE243 MOI 3

| | | | |
|---|---|---|---|
| Mock | 20 | 40 | 30 |
| pREN | 90 | 50 | 50 |
| ORF1 FF (PE243) | 383200 | 375200 | 428400 |
| ORF2 FF (PE243) | 1452000 | 1392000 | 1632000 |
| Main ORF FF (PE243) | 21420000 | 20600000 | 25550000 |

##### Ren Luc

| | | | |
|---|---|---|---|
| Mock | 80 | 60 | 70 |
| pREN | 289900 | 227200 | 317700 |
| ORF1 FF | 191000 | 170700 | 213200 |
| ORF2 FF | 188200 | 151300 | 197700 |
| Main ORF FF | 184800 | 176400 | 222500 |

### Replicate 3

##### FFLuc 24 h p.i.

| | | | | Ren Luc | | | |
|---|---|---|---|---|---|---|---|
| **Mock** | 20 | 20 | 40 | **Mock** | 50 | 100 | 50 |
| **pREN** | 80 | 50 | 90 | **pREN** | 259000 | 244000 | 266000 |
| **ORF1 FF** | 549700 | 335700 | 582200 | **ORF1 FF** | 368500 | 250000 | 417000 |
| **ORF2 FF** | 1887000 | 1136000 | 1855000 | **ORF2 FF** | 369900 | 224800 | 366800 |
| **Main ORF FF** | 38470000 | 26420000 | 42680000 | **Main ORF FF** | 412300 | 301100 | 477400 |

##### FFLuc 24 h p.i. + PE243 MOI 3

| | | | | Ren Luc | | | |
|---|---|---|---|---|---|---|---|
| **Mock** | 20 | 40 | 30 | **Mock** | 80 | 60 | 70 |
| **pREN** | 90 | 50 | 50 | **pREN** | 290000 | 227000 | 318000 |
| **ORF1 FF (PE243)** | 568400 | 589700 | 341700 | **ORF1 FF** | 263800 | 265100 | 191900 |
| **ORF2 FF (PE243)** | 1640000 | 1528000 | 1278000 | **ORF2 FF** | 257400 | 198100 | 178300 |
| **Main ORF FF (PE243)** | 28210000 | 17940000 | 17440000 | **Main ORF FF** | 218700 | 154800 | 171400 |

**Supp Table 2. Percentage of ZIKV E-positive cells in relation to total number of nuclei.**

15 images per virus type at 20X resolution (approx. 400-500 nuclei/image) were quantified for E-positive staining. SDEV represent standard deviation.

| Image number | American WT (% infected cells) | African-like (% infected cells) | uORF1-KO (% infected cells) |
|---|---|---|---|
| **1** | 45.17045455 | 42.30769231 | 7.478632479 |
| **2** | 53.57142857 | 22.23650386 | 7.142857143 |
| **3** | 57.58196721 | 48.00796813 | 10.20408163 |
| **4** | 33.96226415 | 24.37574316 | 10.29810298 |
| **5** | 63.82978723 | 26.24277457 | 7.63546798 |
| **6** | 57.71144279 | 37.5984252 | 4.295942721 |
| **7** | 53.77777778 | 45.56521739 | 6.497175141 |
| **8** | 48.77300613 | 41.84397163 | 7.203389831 |
| **9** | 48.01223242 | 34.24657534 | 2.30125523 |
| **10** | 32.36151603 | 26.05752961 | 7.2319202 |
| **11** | 53.38345865 | 25.79666161 | 10.92592593 |
| **12** | 39.83228512 | 35.66433566 | Bad scanning |
| **13** | 48.0127186 | 35.34482759 | 10.17699115 |
| **14** | 38.60103627 | 41.39534884 | 6.161137441 |
| **15** | 37.55274262 | 32.70348837 | 8.613445378 |
| **Average** | **47.47560787** | **34.62580422** | **7.583308945** |
| **SDEV** | **9.412796729** | **8.238269027** | **2.411265667** |

**Supp Table 3. Percentage of ZIKV E-positive cells that are positive for different cellular markers infected with different ZIKV mutant viruses:**

SDEV indicates standard deviation.

**a. % Cells that are positive for E and nestin**

| Image number | American WT (%) | African-like (%) | uORF1-KO (%) |
|---|---|---|---|
| **1** | 38.9937107 | 35.3535354 | 40 |
| **2** | 33.8461538 | 41.0404624 | 42.4242424 |
| **3** | 36.2989324 | 43.9834025 | 46 |
| **4** | 36.1111111 | 44.3902439 | 44.7368421 |
| **5** | 41.1111111 | 35.2422907 | 45.1612903 |
| **Average** | **37.2722038** | **40.001987** | **43.664475** |
| **SDEV** | 2.81664726 | 4.48468812 | 2.43985954 |

**b. % Cells that are positive for E and MAP2**

| Image number | American WT (%) | African-like (%) | uORF1-KO (%) |
|---|---|---|---|
| **1** | 1.29310345 | 2.09424084 | 0 |
| **2** | 2.47933884 | 1.14503817 | 4.34782609 |
| **3** | 1.25786164 | 0.84745763 | 2.94117647 |
| **4** | 2.5477707 | 1.14285714 | 0 |
| **5** | 3.6036036 | 1.94805195 | 0 |
| **Average** | **2.23633565** | **1.43552914** | **1.45780051** |
| **SDEV** | 0.98394136 | 0.5505567 | 2.05719463 |

**c. % Cells that are positive for E and GFAP**

| Image number | American WT (%) | African-like (%) | uORF1-KO (%) |
|---|---|---|---|
| **1** | 27.6995305 | 31.7647059 | 28.8135593 |
| **2** | 22.6315789 | 28.627451 | N/A |
| **3** | 32.4503311 | 29.2682927 | 36.9565217 |
| **4** | 35.5704698 | 32.0224719 | 42.3076923 |
| **5** | 28.0898876 | 32.8888889 | 31.7073171 |
| **Average** | **29.2883596** | **30.9143621** | **34.9462726** |
| **SDEV** | 4.94308644 | 1.8566943 | 5.95350336 |

**Supp Table 4. Test statistics of ZIKV infection, dissemination and transmission in *Ae. aegypti*.** The table shows the results of the logistic regression analysis. Interactions terms were removed from the final model because their effect was non-significant (*P*<0.05). Blood meal titre was $log_{10}$-transformed.

| | | Infection | | Dissemination | | Transmission | |
|---|---|---|---|---|---|---|---|
| Variable | df | LR $Chi^2$ | P value | LR $Chi^2$ | P value | LR $Chi^2$ | P value |
| Blood meal titre | 1 | 0.3471 | 0.5558 | 0.9425 | 0.3316 | 0.1938 | 0.6598 |
| Experiment | 2 | 1.4035 | 0.4957 | 4.4994 | 0.1054 | 8.8575 | 0.0119 |
| Virus strain | 2 | 4.8015 | 0.0906 | 4.3105 | 0.1159 | 4.9545 | 0.0840 |
| Time | 1 | 0.3726 | 0.5416 | 28.529 | <0.0001 | 6.0300 | 0.0141 |

df: degrees of freedom; LR=likelihood ratio.

#### Supplementary Table 5

##### U251 cells

**American WT (50): African like (50)**

American WT (50%)

| | REP1 | REP2 | REP3 | REP4 | REP5 | REP6 | REP7 | REP8 | Average |
|---|---|---|---|---|---|---|---|---|---|
| **Input 16 h p.i. (p0)** | 50 | | | | | | | | 50 |
| **passage 2 (p2)** | 11.6651 | 25.40687 | 17.15363 | 20.35815 | 8.075772 | 10.56202 | 10.7109 | 10.16782 | 14.2625328 |
| **passage 4 (p4)** | 5.781991 | 22.24511 | 4.610951 | 7.272727 | 4.927007 | 5.607477 | 4.952381 | 5.073801 | 7.55893063 |
| **passage 6 (p6)** | 6.332139 | 18.57143 | 3.435115 | 4.182156 | 8.130841 | 21.31495 | 8.45341 | 11.19617 | 10.2020264 |

African like (50%)

| | REP1 | REP2 | REP3 | REP4 | REP5 | REP6 | REP7 | REP8 | Average |
|---|---|---|---|---|---|---|---|---|---|
| **Input 16 h p.i. (p0)** | 50 | | | | | | | | 50 |
| **passage 2 (p2)** | 88.3349 | 74.59313 | 82.84637 | 79.64185 | 91.92422 | 89.43799 | 89.2891 | 89.83218 | 85.7374675 |
| **passage 4 (p4)** | 94.21801 | 77.75489 | 95.38905 | 92.72727 | 95.07299 | 94.39252 | 95.04762 | 94.9262 | 92.4410688 |
| **passage 6 (p6)** | 93.66786 | 81.42857 | 96.56489 | 95.81784 | 91.86916 | 78.68505 | 91.54659 | 88.80383 | 89.7979738 |

**American WT (90): African like (10)**

American WT (90%)

| | REP1 | REP2 | REP3 | REP4 | REP5 | REP6 | REP7 | REP8 | Average |
|---|---|---|---|---|---|---|---|---|---|
| **Input 16 h p.i. (p0** | 90 | | | | | | | | 90 |
| **passage 2 (p2)** | 48.74031 | 63.56757 | 56.3745 | 58.77712 | 43.35472 | 32.90441 | 36.13139 | 43.41988 | 47.9087375 |
| **passage 4 (p4)** | 42.82655 | 28.26923 | 19.70121 | 17.65258 | 19.1022 | 8.593012 | 11.84702 | 14.94253 | 20.3667915 |
| **passage 6 (p6)** | 14.30063 | 19.2053 | 15.15435 | 10.06464 | 15.06977 | 8.211143 | 11.95551 | 11.29477 | 13.1570141 |

African like (10%)

| | REP1 | REP2 | REP3 | REP4 | REP5 | REP6 | REP7 | REP8 | Average |
|---|---|---|---|---|---|---|---|---|---|
| **Input 16 h p.i. (p0** | 10 | | | | | | | | 10 |
| **passage 2 (p2)** | 51.25969 | 36.43243 | 43.6255 | 41.22288 | 56.64528 | 67.09559 | 63.86861 | 56.58012 | 52.0912625 |
| **passage 4 (p4)** | 57.17345 | 71.73077 | 80.29879 | 82.34742 | 80.8978 | 91.40699 | 88.15298 | 85.05747 | 79.6332088 |
| **passage 6 (p6)** | 85.69937 | 80.7947 | 84.84565 | 89.93536 | 84.93023 | 91.78886 | 88.04449 | 88.70523 | 86.8429863 |

**American WT (50): uORF1-KO (50)**

American WT (50%)

| | REP1 | REP2 | REP3 | REP4 | REP5 | REP6 | REP7 | REP8 | Average |
|---|---|---|---|---|---|---|---|---|---|
| **Input 16 h p.i. (p0)** | 50 | | | | | | | | 50 |
| **passage 2 (p2)** | 18.93584 | 59.53846 | 39.68254 | 46.19799 | 30.61516 | 36.88639 | 34.64912 | 35.38462 | 37.736265 |
| **passage 4 (p4)** | 29.82172 | 46.12188 | *N/A* | 23.71428 | 22.02073 | 19.81004 | 16.71642 | 30.31989 | 26.9321371 |
| **passage 6 (p6)** | 32.80507 | 39.85611 | 16.71512 | 31.4456 | 12.1118 | 7.153729 | 15.18152 | 37.61194 | 24.1101111 |

uORF1-KO (50%)

| | REP1 | REP2 | REP3 | REP4 | REP5 | REP6 | REP7 | REP8 | Average |
|---|---|---|---|---|---|---|---|---|---|
| **Input 16 h p.i. (p0)** | 50 | | | | | | | | 50 |
| **passage 2 (p2)** | 81.06416 | 40.46154 | 60.31746 | 53.80201 | 69.38483 | 63.11361 | 65.35088 | 64.61539 | 62.263735 |
| **passage 4 (p4)** | 70.17828 | 53.87812 | *N/A* | 76.28571 | 77.97927 | 80.18996 | 83.28358 | 69.68011 | 73.0678614 |
| **passage 6 (p6)** | 67.19493 | 60.14389 | 83.28488 | 68.5544 | 87.8882 | 92.84627 | 84.81848 | 62.38806 | 75.8898888 |

**American WT (90): uORF1-KO (10)**

American WT (90%)

| | REP1 | REP2 | REP3 | REP4 | REP5 | REP6 | REP7 | REP8 | Average |
|---|---|---|---|---|---|---|---|---|---|
| **Input 16 h p.i. (p0** | 90 | | | | | | | | 90 |
| **passage 2 (p2)** | 63.06306 | 66.59836 | 65.16129 | 68.31832 | 68.13509 | 68.07512 | 74.09638 | 67.44526 | 67.61161 |
| **passage 4 (p4)** | 2.335165 | 5.443235 | 5.604203 | 5.28109 | 1.618123 | 9.204369 | 6.445993 | 2.056075 | 4.74853163 |
| **passage 6 (p6)** | 1.186441 | 0 | 1.083591 | 0 | 5.28169 | 0.1972387 | 0.525394 | 0 | 1.03429434 |

uORF1-KO (10%)

| | REP1 | REP2 | REP3 | REP4 | REP5 | REP6 | REP7 | REP8 | Average |
|---|---|---|---|---|---|---|---|---|---|
| **Input 16 h p.i. (p0** | 10 | | | | | | | | 10 |
| **passage 2 (p2)** | 36.93694 | 33.40164 | 34.83871 | 31.68168 | 31.8649 | 31.92488 | 25.90361 | 32.55474 | 32.3883875 |
| **passage 4 (p4)** | 97.66483 | 94.55676 | 94.3958 | 94.71891 | 98.38187 | 90.79563 | 93.55401 | 97.94392 | 95.2514663 |
| **passage 6 (p6)** | 98.81356 | 100 | 98.91641 | 100 | 94.71831 | 99.80276 | 99.47461 | 100 | 98.9657063 |

**American WT (50): uORF2-PTC1 (50)**

American WT (50%)

| | REP1 | REP2 | REP3 | REP4 | REP5 | REP6 | REP7 | REP8 | Average |
|---|---|---|---|---|---|---|---|---|---|
| **Input 16 h p.i. (p0)** | 50 | | | | | | | | 50 |
| **passage 2 (p2)** | 54.02299 | 23.91681 | 20.91603 | 20.95532 | 19.97106 | 17.27141 | 25.82973 | 19.30556 | 25.2736138 |
| **passage 4 (p4)** | 37.92614 | 31.40741 | 32.95454 | 23.9645 | 20.17291 | 17.27528 | 19.69925 | 22.15827 | 25.6947875 |
| **passage 6 (p6)** | 75.87719 | 58.2404 | 70.97345 | 34.24069 | 44.91525 | 23.11321 | 32.38381 | 35.20115 | 46.8681438 |

uORF2-PTC1 (50%)

| | REP1 | REP2 | REP3 | REP4 | REP5 | REP6 | REP7 | REP8 | Average |
|---|---|---|---|---|---|---|---|---|---|
| **Input 16 h p.i. (p0)** | 50 | | | | | | | | 50 |
| **passage 2 (p2)** | 45.97701 | 76.08319 | 79.08397 | 79.04469 | 80.02895 | 82.72859 | 74.17027 | 80.69444 | 74.7263888 |
| **passage 4 (p4)** | 62.07386 | 68.59259 | 67.04546 | 76.0355 | 79.82709 | 82.72472 | 80.30075 | 77.84173 | 74.3052125 |
| **passage 6 (p6)** | 24.12281 | 41.7596 | 29.02655 | 65.75932 | 55.08475 | 76.8868 | 67.6162 | 64.79885 | 53.13186 |

**American WT (10): uORF2-PTC1 (90)**

American WT (10%)

| | REP1 | REP2 | REP3 | REP4 | REP5 | REP6 | REP7 | REP8 | Average |
|---|---|---|---|---|---|---|---|---|---|
| **Input 16 h p.i. (p0** | 10 | | | | | | | | 10 |
| **passage 2 (p2)** | 16.66667 | 44.51613 | 21.71533 | 15.478 | 7.608696 | 5.648855 | 5.279035 | 4.431314 | 15.1680038 |
| **passage 4 (p4)** | 8.544304 | 5.263158 | 3.614458 | 5.325444 | 3.385049 | 5.738881 | 5.028736 | 5.192878 | 5.2616135 |
| **passage 6 (p6)** | 2.72 | 5.279503 | 3.994294 | 4.069767 | 5.961252 | 9.563994 | 5.675147 | 10.40268 | 5.95832963 |

uORF2-PTC1 (90%)

| | REP1 | REP2 | REP3 | REP4 | REP5 | REP6 | REP7 | REP8 | Average |
|---|---|---|---|---|---|---|---|---|---|
| **Input 16 h p.i. (p0** | 90 | | | | | | | | 90 |
| **passage 2 (p2)** | 83.33334 | 55.48387 | 78.28467 | 84.522 | 92.3913 | 94.35114 | 94.72096 | 95.56869 | 84.8319963 |
| **passage 4 (p4)** | 91.4557 | 94.73684 | 96.38554 | 94.67455 | 96.61495 | 94.26112 | 94.97127 | 94.80712 | 94.7383863 |
| **passage 6 (p6)** | 97.28 | 94.7205 | 96.00571 | 95.93023 | 94.03875 | 90.436 | 94.32485 | 89.59731 | 94.0416688 |

##### Vero cells

**American WT (50): African like (50)**

American WT (50%)

| | REP1 | REP2 | REP3 | REP4 | REP5 | REP6 | REP7 | REP8 | Average |
|---|---|---|---|---|---|---|---|---|---|
| **Input 16 h p.i. (p0)** | 50 | | | | | | | | 50 |
| **passage 2 (p2)** | 26.23427 | 35.80586 | 39.50504 | 32.56039 | 39.83441 | 27.49546 | 28.44677 | 27.95415 | 32.2295438 |

| | | | | | | | | | |
|---|---|---|---|---|---|---|---|---|---|
| **passage 4 (p4)** | 21.87812 | 18.62653 | 25.39289 | 27.0032 | 27.71203 | 24.73532 | 25.13612 | 26.9697 | 24.6817388 |
| **passage 6 (p6)** | 23.79249 | 11.50943 | 11.39122 | 10.6383 | 17.56757 | 15.72505 | 17.54717 | 15.67308 | 15.4805388 |

**African like (50%)**

| | REP1 | REP2 | REP3 | REP4 | REP5 | REP6 | REP7 | REP8 | Average |
|---|---|---|---|---|---|---|---|---|---|
| **Input 16 h p.i. (p0)** | 50 | | | | | | | | 50 |
| **passage 2 (p2)** | 73.76573 | 64.19414 | 60.49496 | 67.43961 | 60.16559 | 72.50454 | 71.55323 | 72.04585 | 67.7704563 |
| **passage 4 (p4)** | 78.12188 | 81.37347 | 74.60712 | 72.9968 | 72.28797 | 75.26468 | 74.86388 | 73.0303 | 75.3182625 |
| **passage 6 (p6)** | 76.20751 | 88.49056 | 88.60878 | 89.3617 | 82.43243 | 84.27496 | 82.45283 | 84.32692 | 84.5194613 |

**American WT (50): uORF1-KO (50)**

**American WT (50%)**

| | REP1 | REP2 | REP3 | REP4 | REP5 | REP6 | REP7 | REP8 | Average |
|---|---|---|---|---|---|---|---|---|---|
| **Input 16 h p.i. (p0)** | 50 | | | | | | | | 50 |
| **passage 2 (p2)** | 42.54777 | 42.83361 | 45.8671 | 46.60574 | 48.91641 | 49.44 | 48 | 48.76033 | 46.62137 |
| **passage 4 (p4)** | 51.78849 | 27.61506 | 35.79137 | 48.49355 | 46.98413 | 53.03644 | 41.36213 | 45.42936 | 43.8125663 |
| **passage 6 (p6)** | 33.95683 | 19.54198 | 23.68421 | 31.62518 | 45.8886 | 50.14045 | 39.36022 | 46.37483 | 36.3215375 |

**uORF1-KO (50%)**

| | REP1 | REP2 | REP3 | REP4 | REP5 | REP6 | REP7 | REP8 | Average |
|---|---|---|---|---|---|---|---|---|---|
| **Input 16 h p.i. (p0)** | 50 | | | | | | | | 50 |
| **passage 2 (p2)** | 57.45223 | 57.16639 | 54.1329 | 53.39426 | 51.08359 | 50.56 | 52 | 51.23967 | 53.37863 |
| **passage 4 (p4)** | 48.21151 | 72.38493 | 64.20863 | 51.50645 | 53.01587 | 46.96356 | 58.63787 | 54.57064 | 56.1874325 |
| **passage 6 (p6)** | 66.04317 | 80.45802 | 76.31579 | 68.37482 | 54.1114 | 49.85955 | 60.63978 | 53.62517 | 63.6784625 |

**American WT (50): uORF2-PTC1 (50)**

**American WT (50%)**

| | REP1 | REP2 | REP3 | REP4 | REP5 | REP6 | REP7 | REP8 | Average |
|---|---|---|---|---|---|---|---|---|---|
| **Input 16 h p.i. (p0)** | 50 | | | | | | | | 50 |
| **passage 2 (p2)** | 67.0469 | 59.23077 | 62.41331 | 71.24464 | 58.32241 | 59.86577 | 61.34021 | 62.21374 | 62.7097188 |
| **passage 4 (p4)** | 80.28503 | 73.10513 | 73.38618 | 69.83051 | 59.42928 | 67.42712 | 68.9441 | 73.14487 | 70.6940275 |
| **passage 6 (p6)** | ***N/A*** | 74.0227 | 70 | 75.99039 | 70.51282 | 72.40224 | 76.17952 | 65.19139 | 72.0427229 |

**uORF2-PTC1 (50%)**

| | REP1 | REP2 | REP3 | REP4 | REP5 | REP6 | REP7 | REP8 | Average |
|---|---|---|---|---|---|---|---|---|---|
| **Input 16 h p.i. (p0)** | 50 | | | | | | | | 50 |
| **passage 2 (p2)** | 32.95311 | 40.76923 | 37.58669 | 28.75537 | 41.67759 | 40.13423 | 38.65979 | 37.78626 | 37.2902838 |
| **passage 4 (p4)** | 19.71496 | 26.89487 | 26.61382 | 30.16949 | 40.57072 | 32.57288 | 31.0559 | 26.85512 | 29.30597 |
| **passage 6 (p6)** | ***N/A*** | 25.9773 | 30 | 24.0096 | 29.48718 | 27.59776 | 23.82048 | 34.80861 | 27.9572757 |

#### Supp Table 6: Quantification of the collapse of vimentin in infected cells:

**Key legend for the collapse of vimentin:**

**1.** No collapse of vimentin
**2.** Perinuclear accumulation of vimentin
**3.** Starting of the formation of the vimentin dot
**4.** Complete vimentin dot formation

**Experiment 1:**

| | Collapse of vimentin (AU) (nº of cells) | | | | Total nº of cells |
|---|---|---|---|---|---|
| **Sample** | **1** | **2** | **3** | **4** | |
| American WT (18 h p.i.) | 76 | 65 | 18 | 2 | 161 |
| uORF1-KO (18 h p.i.) | 83 | 55 | 4 | 3 | 146 |
| American WT (24 h p.i.) | 19 | 46 | 64 | 59 | 188 |
| uORF1-KO (24 h p.i.) | 29 | 78 | 23 | 19 | 149 |

**Experiment 2:**

| | Collapse of vimentin (AU) (nº of cells) | | | | Total nº of cells |
|---|---|---|---|---|---|
| **Sample** | **1** | **2** | **3** | **4** | |
| American WT (18 h p.i.) | 51 | 72 | 34 | 2 | 159 |
| uORF1-KO (18 h p.i.) | 55 | 39 | 15 | 2 | 111 |
| American WT (24 h p.i.) | 11 | 59 | 73 | 39 | 182 |
| uORF1-KO (24 h p.i.) | 32 | 74 | 52 | 13 | 171 |

**Supplementary Table 7. List of oligonucleotides used.** Note *fwd* indicates forward primer, and *rev* indicates reverse primer.

| **Name** | **Sequence (5′-3′)** |
|---|---|
| *uORF1 KO, fwd* | TGTGTGAATCAGACTACGACAGTTCGAGTTT |
| *uORF1 KO, rev* | AAACTCGAACTGTCGTAGTCTGATTCACACA |
| *African ORF like, fwd* | CAGTATCAACAGGTTTAATTTGGATTTGGAAACGAGA |
| *African ORF like, rev* | TCTCGTTTCCAAATCCAAATTAAACCTGTTGATACTG |
| *uORF2 KO, fwd* | AACAGGTTTTATTTAGATTTGGAAACGAGA |
| *uORF2 KO, rev* | TCTCGTTTCCAAATCTAAAATAAAACCTGTT |
| *uORF2-AUG, fwd* | AACAGGTTTTATTATGGATTTGGAAACGAGA |
| *uORF2-AUG, rev* | TCTCGTTTCCAAATCCATAATAAAACCTGTT |
| *uORF2-PTC1, fwd* | TTGGATTTGGAAACGTGAGTTTCTGGTCATG |
| *uORF2-PTC1, rev* | CATGACCAGAAACTCACGTTTCCAAATCCAA |
| *uORF1 in frame pSGD, fwd* | GCGCCTCGAGATAAGTTGTTGATCTGTGTGAATCAG AC |
| *uORF1 in frame pSGD, rev* | GCGCAGATCTATTTCTTTTTTGGGGTTTTCCATGACC AG |
| *uORF2 in frame pSGD, fwd* | GCGCCTCGAGATAAGTTGTTGATCTGTGTGAATCAG AC |
| *uORF2 in frame pSGD, rev* | GCGCAGATCTAAGCCCCCAAAGGGGCTCACACGGG |
| *Main ORF in frame pSGD, fwd* | GCGCCTCGAGATAAGTTGTTGATCTGTGTGAATCAG AC |
| *Main ORF in frame pSGD, rev* | GCGCAGATCTGCCCCCAAAGGGGCTCACACGGGCTA CTCCG |
| *T7 upstream of ZIKV uORFs, fwd* | GACTCACTATAGGGAGTTGTTGATCTGTGTGAATC |
| *T7 upstream of ZIKV uORFs, rev* | GATTCACACAGATCAACAACTCCCTATAGTGAGTC |
| *uORF2-PTC34, fwd* | CGGAGTAGCCCGTGTTAGCCCCTTTGGGGGC |
| *uORF2-PTC34, rev* | GCCCCCAAAGGGGCTAACACGGGCTACTCCG |
| *uORF2-FS1, fwd* | CGAGAGTTTCTGGTCTTAGAAAACCCAAAAAAGA |
| *uORF2-FS1, rev* | TCTTTTTTGGGTTTTCTAAGACCAGAAACTCTCG |
| *uORF2-FS2, fwd* | GTTTCTGGTCATGAAGAACCCAAAAAAGAAA |
| *uORF2-FS2, rev* | TTTCTTTTTTGGGTTCTTCATGACCAGAAAC |
| *uORF2-FS3, fwd* | GGTCATGAAAAACCCGAAGAAGAAATCCGGAGGA |
| *uORF2-FS3, rev* | TCCTCCGGATTTCTTCTTCGGGTTTTTCATGACC |
| *uORF2-alternative initiation Mut1, fwd* | AAACCCAAA AAAGAAGTCCGGAGGATTCCGG |
| *uORF2-alternative initiation Mut1, rev* | CCGGAATCCTCCGGACTTCTTTTTTGGGTTT |
| *uORF2-alternative initiation Mut2, fwd* | A AAAGAAGTCCGGAGAATTCCGGATTGTCAA |
| *uORF2-alternative initiation Mut2, rev* | TTGACAATCCGGAATTCTCCGGACTTCTTTT |
| *uORF2-alternative initiation Mut3, fwd* | AAAGAAGTCCGGAGAGTTCCGGATTGTCAA T |
| *uORF2-alternative initiation Mut3, rev* | ATTGACAATCCGGAACTCTCCGGACTTCTTT |
| *uORF2-alternative initiation Mut4,* | TCAATATGCTAAAACACGGAGTAGCCCGTGT |

| | |
|---|---|
| *fwd* | |
| *uORF2-alternative initiation Mut4, rev* | ACACGGGCTACTCCGTGTTTTAGCATATTGA |
| *uORF1-TAP in pCAG, fwd* | GCGCTTAATTAAACCATGCGACAGTTCGAGTTTGAAGC |
| *uORF1-TAP in pCAG, rev* | GCGCCTTAAGTTACTTGTCATCGTCATCCTTG |
| *uORF1-mCherry in pCAG (PCR1), fwd* | GCGCTTAATTAAACCATGCGACAGTTCGAGTTTGAAGCG |
| *uORF1-mCherry in pCAG (PCR1), rev* | GTTATCCTCCTCGCCCTTGCTCACGGCTGATGACCAGAAACTCTCGTTTCCAAA |
| *uORF1-mCherry in pCAG (PCR2), fwd* | TTTGGAAACGAGAGTTTCTGGTCATCAGCCGTGAGCAAGGGCGAGGAGGATAAC |
| *uORF1-mCherry in pCAG (PCR2), rev* | GCGCCTTAAGTTACTTGTACAGCTCGTCCATGCCGCC |
| *uORF2-FLAG in pCAG, fwd* | GCGCTTAATTAAACCATGGATTTGGAAACGAGAGTTTC |
| *uORF2-FLAG in pCAG, rev* | GCGCCTTAAGCTACTTGTCGTCATCGTCTTTGTAGTCTTGATGAGACCCAGTGATGGC |
| *African ORF-FLAG in pCAG, fwd* | GCGCTTAATTAAACCATGCGACAGTTCGAGTTTGAAGC |
| *African ORF-FLAG in pCAG, rev* | GCGCCTTAAGCTACTTGTCGTCATCGTCTTTGTAGTCTTGATGAGACCCAGTGATGGC |
| *SHAPE analysis, fwd* | GGCTACTCCGCGTTTTAGCATATTG |
| *SHAPE analysis, rev* | TCTTCAAGCCCCCAAAGGGGCTCAC |
| *To amplify ZIKV 5' UTR, fwd* | GTTGTTGATCTGTGTGAATCAG |
| *To amplify ZIKV 5' UTR, rev* | TATTGATGAGACCCAGTGATGGC |
| *To sequence ZIKV 5' UTR, rev* | GACCCAGCAGAAGTCCGGCTGGC |

4

**Supplementary Table 8. List of antibodies used.**

#### - Immunoblotting

| Antibody | Host | Type | Source | Dilution |
|---|---|---|---|---|
| | | **Primary Antibodies** | | |
| Anti-GAPDH | Mouse | IgM | Sigma-Aldrich, G8795 | 1:20,000 |
| Anti-FLAG | Mouse | IgG | Sigma-Aldrich, SAB4301135 | 1:2,000 |
| Anti-E protein | Rabbit | IgG | GeneTex, GTX133314 | 1:1,000 |
| Anti-Lamin A+C | Rabbit | IgG | Abcam, ab108922 | 1:1,000 |
| Anti-H2A | Rabbit | IgG | Abcam, ab1777308 | 1:1,000 |
| Anti-Vimentin | Mouse | IgG1 | Abcam, ab8069 | 1:1,000 |
| Anti-ERp72 | Rabbit | IgG | Cell Signaling, 5033 | 1:1,000 |
| Anti-eIF2α | Rabbit | IgG | Cell Signaling, 9722 | 1:1,000 |
| Anti-p-eIF2α | Rabbit | IgG | Cell Signaling, 9721 | 1:1,000 |
| Anti-mCherry | Rabbit | IgG | Abcam, ab167453 | 1:1,000 |
| | | **Secondary Antibodies** | | |
| Anti-Rabbit | Goat | 800 | Licor, IRDye 926-32211 | 1:1,000 |
| Anti-Mouse IgM | Donkey | 680 | Licor, IRDye 926-68180 | 1:1,000 |
| Anti-Mouse | Goat | 800 | Licor, IRDye 926-32210 | 1:1,000 |

#### - Immunofluorescence

| Antibody | Host | Type | Source, Catalogue # | Dilution |
|---|---|---|---|---|
| | | **Primary Antibodies** | | |
| Anti-E protein | Rabbit | IgG | GeneTex, GTX133314 | 1:200 |
| Anti-flavivirus Ag group | Mouse | IgG | Merck, D1-4G2-4-15 | 1:200 |
| Anti-flavivirus Ag group | Rabbit | IgG | Absolute antibody, Ab00230-23.0 | 1:200 |
| Anti-FLAG | Mouse | IgG | Sigma-Aldrich, SAB4301135 | 1:500 |
| Anti-Vimentin | Mouse | IgG1 | Abcam, ab8069 | 1:500 |
| Anti-Actin | Mouse | IgG | Proteintech, 66009 | 1:500 |
| Anti-Tubulin | Rat | Hybridoma | Kind gift from Dr Colin Crump | 1:10 |
| Anti-MAP2 | Goat | IgG | antibodies.com, A104327 | 1:1,000 |
| Anti-Nestin | Mouse | IgG1 | Abcam, ab22035 | 1:500 |

| | | | | |
|---|---|---|---|---|
| Anti-GFAP | Rabbit | IgG | antibodies.com, A85419 | 1:1,000 |
| | | **Secondary Antibodies** | | |
| Anti-Rabbit | Goat | 488 | Invitrogen Alexa Fluor, A11008 | 1:1,000 |
| Anti-Rabbit | Donkey | 488 | Abcam, ab150073 | 1:1,000 |
| Anti-Mouse | Goat | 488 | Invitrogen Alexa Fluor, A11001 | 1:1,000 |
| Anti- Rabbit | Donkey | 594 | Invitrogen Alexa Fluor, A21207 | 1:1,000 |
| Anti-Mouse | Donkey | 594 | Invitrogen Alexa Fluor, A21203 | 1:1,000 |
| Anti-Rat | Goat | 568 | Invitrogen Alexa Fluor, A11077 | 1:1,000 |

**Supp Table 9. Genbank accession numbers for rRNA**

| | **rRNA accession numbers** |
|---|---|
| ***Chlorocebus sabaeus*** | NR_003287.2, NR_023379.1, NR_003285.2, NR_003286.2, AY603036.1, AF420058.1, AF420040.1, AY633510.1, AF352382.1, L35185.1, DQ983926.1, KJ193255.1, M30951.1, M30950.1, M30952.1, KJ193272.1, KJ193259.1, KJ193258.1, KJ193256.1, KJ193255.1, KJ193045.1, KJ193042.1, KJ193044.1, KJ193041.1, KJ193019.1, KJ193018.1, KJ193017.1, AF420040.1 |
| ***Homo sapiens*** | NR_003287.4, NR_023379.1, NR_003285.3 and NR_003286.4 |
| ***Aedes albopictus*** | L22060, DQ397934.1, DQ397935.1, JX522172.1, AB085210.1, X57172.1, XR_003895909.1 |

**Supp Table 10. Designated regions to calculate the number of reads per phase**

<table>
<tr><th>Virus</th><th>Region of interest</th><th>Coordinates barcharts</th></tr>
<tr><td rowspan="3">PE243, mutant viruses (American WT, uORF1-KO, uORF2-PTC1)</td><td>uORF1</td><td>25-79</td></tr>
<tr><td>uORF2</td><td>80-107</td></tr>
<tr><td>Main ORF</td><td>311-480</td></tr>
<tr><td rowspan="2">Dak84, mutant virus (African-like)</td><td>Afr uORF</td><td>25-106</td></tr>
<tr><td>Main ORF</td><td>311-480</td></tr>
</table>
